# Dynamic Co-Modulation (DyCoM): A Unified Operator Framework for Dynamic Connectivity in Neuroimaging

**DOI:** 10.64898/2026.01.22.701132

**Authors:** Sir-Lord Wiafe, Najme Soleimani, Armin Iraji, Tülay Adali, Vince D. Calhoun

## Abstract

Dynamic connectivity is central to understanding time-varying interactions between brain regions. Despite decades of methodological development, approaches to measuring dynamic connectivity remain fragmented, leading to inconsistent findings, limited comparability across studies, and difficulty attributing observed effects to computational choices. Here, we introduce dynamic co-modulation (DyCoM), a compact operator-level framework that expresses dynamic connectivity estimators as compositions of a small set of fundamental signal processing operations. Using simulations and resting-state fMRI data, we show that DyCoM disentangles previously conflated findings by revealing that lower-order sensory and higher-order executive control neurobiological signatures, state-transition sensitivity, and medication-linked clinical associations arise from distinct operator choices within a single unified framework. Together, these results establish DyCoM as a unifying foundation for dynamic interaction analysis, revealing how differences in estimator design give rise to divergent biological interpretations and offering a principled and extensible framework for coherence, interpretability, and estimator development.

## 1. INTRODUCTION

Time-resolved functional connectivity, often referred to as dynamic connectivity, has become a central tool for studying nonstationary interactions in fMRI data^1,2^. A wide range of dynamic connectivity estimators have been proposed, including sliding window Pearson correlation (SWPC)^3^, edge-based time series^4^, filter bank and multiscale approaches^5^, and amplitude- or phase-based measures^6,7^. These methods have been used to characterize transient network states, disease-related alterations, and cognitive variability across time.

However, the proliferation of dynamic connectivity estimators, often introduced as distinct analytical approaches, has resulted in a landscape where empirical conclusions depend on methodological choices that are rarely made explicit or interpreted mechanistically, contributing to inconsistent findings across studies. Although some general frameworks for functional connectivity have been proposed^8,9^, these efforts primarily organize existing methods after the fact, grouping them by what they measure or by their analysis goals, rather than showing how the methods themselves are built from a common set of operations. For example, the recent DySCo framework organizes dynamic connectivity measures around a common recurrence-based formalism for comparison purposes^9^; DyCoM differs in being generative at the operator level, specifying the computational components from which estimators are constructed, so that existing methods are recovered and new ones derived from the same building blocks. Consequently, relationships between estimators are typically established retrospectively, and new methods require ad hoc justification instead of arising naturally from a shared generative structure. This obscures the sources of empirical discrepancies and hinders the accumulation of coherent theory, limiting systematic comparison and principled methodological development.

For example, although many studies nominally employ sliding window correlation or related dynamic connectivity estimators, their implementations differ in critical ways, including windowing strategy, normalization, signal representation, and temporal integration, often without clear justification or standardization^10^. Even subtle differences in these choices can substantially alter connectivity estimates and their biological interpretation^11^. In schizophrenia, dynamic connectivity studies have emphasized different network-level abnormalities, including heightened sensory and visual network engagement^12^, fronto-temporal dysconnectivity^13^, and executive control disruptions^14^, sometimes within comparable clinical populations. Although these findings are not inherently contradictory, the dominant conclusions regarding which systems are most disrupted vary across studies, raising the possibility that methodological differences shape which effects are detected or emphasized. Moreover, several dynamic estimators show similar associations with cognitive measures, such as processing speed, and, in some cases, yield highly overlapping behavioral and clinical profiles^15^. Without a principled framework and operator-level interpretability, it remains unclear whether divergent findings reflect genuinely distinct neurobiological processes or whether apparent agreement and reproducibility arise from shared computational structure and estimator redundancy^15,16^.

From a signal processing perspective, many dynamic connectivity estimators can be interpreted through the lens of modulation^17^. In classical communication theory, modulation describes how information is conveyed by varying properties of a signal, such as amplitude, phase, or frequency, relative to a reference waveform^18^. In neuroscience, however, there is no clear distinction between reference and information-bearing signals. Instead, interacting neural time series mutually influence each other’s fluctuations over time. This motivates the concept of “*co-modulation*”, in which two signals jointly shape each other’s amplitude, phase, or frequency content over time. Under this interpretation, correlation-based connectivity reflects amplitude co-modulation^17,19^, while phase synchrony and related measures reflect phase co-modulation^17,20^. The term co-fluctuation is often used in the fMRI literature to denote the instantaneous product of standardized time series, emphasizing temporal deviation as the defining feature of interaction^4,21^. However, at each time point, the outer product of regional signals forms a rank-one interaction matrix, a matrix of all pairwise products generated by a single shared activity pattern, that reflects coordinated modulation across regions or networks. Framing interaction as co-modulation shifts the emphasis from deviation over time to structured joint expression across systems and connects neuroimaging analyses to an established body of signal processing theory built on products of signal pairs, known as bilinear analysis.

Among the different forms of co-modulation, amplitude-based co-modulation has been implicitly advanced through the growing use of edge-based time series, where interactions between pairs of regions are represented directly at each time point^4^. Instantaneous correlation, defined as the pointwise product of two standardized signals, provides a canonical example of this approach^22^. Importantly, this bilinear interaction is not a novel construct, but rather a direct reappearance of classical bilinear energy formulations that have long been studied in time-frequency analysis. In particular, the pointwise product corresponds to the zero-lag, time-domain component of bilinear distributions such as the cross Wigner-Ville representation, a classical representation describing how the joint energy of two signals distributes over time and frequency^23,24^. From this perspective, edge-based time series can be viewed as a special case of these earlier bilinear frameworks, retaining only the same-time interaction between the two signals. Despite its apparent simplicity, this instantaneous bilinear form serves as a fundamental building block from which many dynamic connectivity estimators are constructed, either implicitly or explicitly, through subsequent filtering, normalization, or aggregation steps.

Although many dynamic connectivity estimators rely on closely related instantaneous interaction terms, they are rarely presented within a common mathematical structure. Sliding window correlation, for example, integrates instantaneous products over time while estimating local means and variances for normalization^3^. Filter bank approaches extend this idea by decomposing interaction trajectories across frequency bands^5^. Adaptive and recursive formulations rely on local representations of the data, implicitly reducing slow drifts and shared variance^25^, and phase and amplitude-envelope methods apply analogous operations after nonlinear signal transformations^6,7^. Despite these shared computational elements, such approaches are typically developed and implemented as separate pipelines, with key design choices embedded implicitly rather than expressed through a unified operator form. As a result, similarities between methods are obscured at the implementation level, and methodological differences are often attributed to surface distinctions rather than to specific computational operators.

Here, we introduce dynamic co-modulation (DyCoM), a unified operator framework for time-resolved connectivity analysis. DyCoM expresses dynamic connectivity as an ordered composition of four modular operations acting on pairs of signals: a representational transformation, an instantaneous interaction or energy mapping, temporal integration across a chosen timescale, and optional normalization. Rather than defining connectivity through a single estimator, DyCoM formalizes dynamic connectivity as a family of estimators generated by specific parameterizations of these operators. This operator-level decomposition reveals that many widely used methods differ not in their fundamental computational logic, but in where and how these operations are instantiated. This provides a common generative structure that links existing approaches, supports principled method development, and allows method-dependent findings to be traced to specific computational choices.

While the DyCoM framework naturally accommodates amplitude-, phase-, and frequency-based co-modulation, the present study focuses on amplitude co-modulation corresponding to correlation-based dynamic connectivity. This focus reflects the widespread use of correlation-based estimators in fMRI and their central role in contemporary dynamic functional connectivity analyses^1^. Importantly, DyCoM does not restrict the form of instantaneous interaction or temporal integration, allowing windowed, windowless, adaptive, and multiscale estimators to be expressed within the same operator framework. Phase- and frequency-based instantiations arise from alternative representational and interaction operators and are fully compatible with the proposed formulation, although they are not examined empirically here.

Within this scope, we develop DyCoM as a unified operator framework that exposes the shared computational structure underlying widely used amplitude-based dynamic connectivity estimators. The framework makes explicit how common methods arise from specific choices of representation, instantaneous interaction, temporal integration, and normalization, and it provides a constructive pathway for extending these components in a principled manner. As a concrete demonstration of this generative capability, we derive a new estimator that instantiates all four DyCoM operators within a single formulation. We validate DyCoM using simulations and resting-state fMRI data from individuals with schizophrenia and healthy controls, demonstrating that specific operator choices systematically produce distinct patterns of dynamic connectivity across cognitive and clinical dimensions. By making these computational pathways explicit, DyCoM reveals how diverse connectivity signatures can emerge from a common generative structure. As a shared computational language, it unifies fragmented methodologies, enabling coherent and interpretable progress in dynamic connectivity research.

## 2. METHODS

### 2.1. General framework

Dynamic co-modulation quantifies how multivariate time-varying system components interact through moment-to-moment co-fluctuations across defined temporal scales. Instead of relying on a fixed estimator, DyCoM expresses dynamic connectivity as a sequence of four modular operators: representation, instantaneous energy, temporal integration, and normalization. Together, these operators generate a family of dynamic connectivity measures. This modular decomposition yields a unified mathematical framework that recovers existing estimators and enables principled construction of new ones. Figure 1 provides a schematic overview of the DyCoM pipeline, illustrating how these four operators are composed and how common dynamic connectivity estimators arise from specific operator choices. The figure is intended as a conceptual guide to the framework rather than a detailed specification of any single method. Formal definitions of each operator and their parameterizations are introduced in the sections that follow.

**Figure 1:**
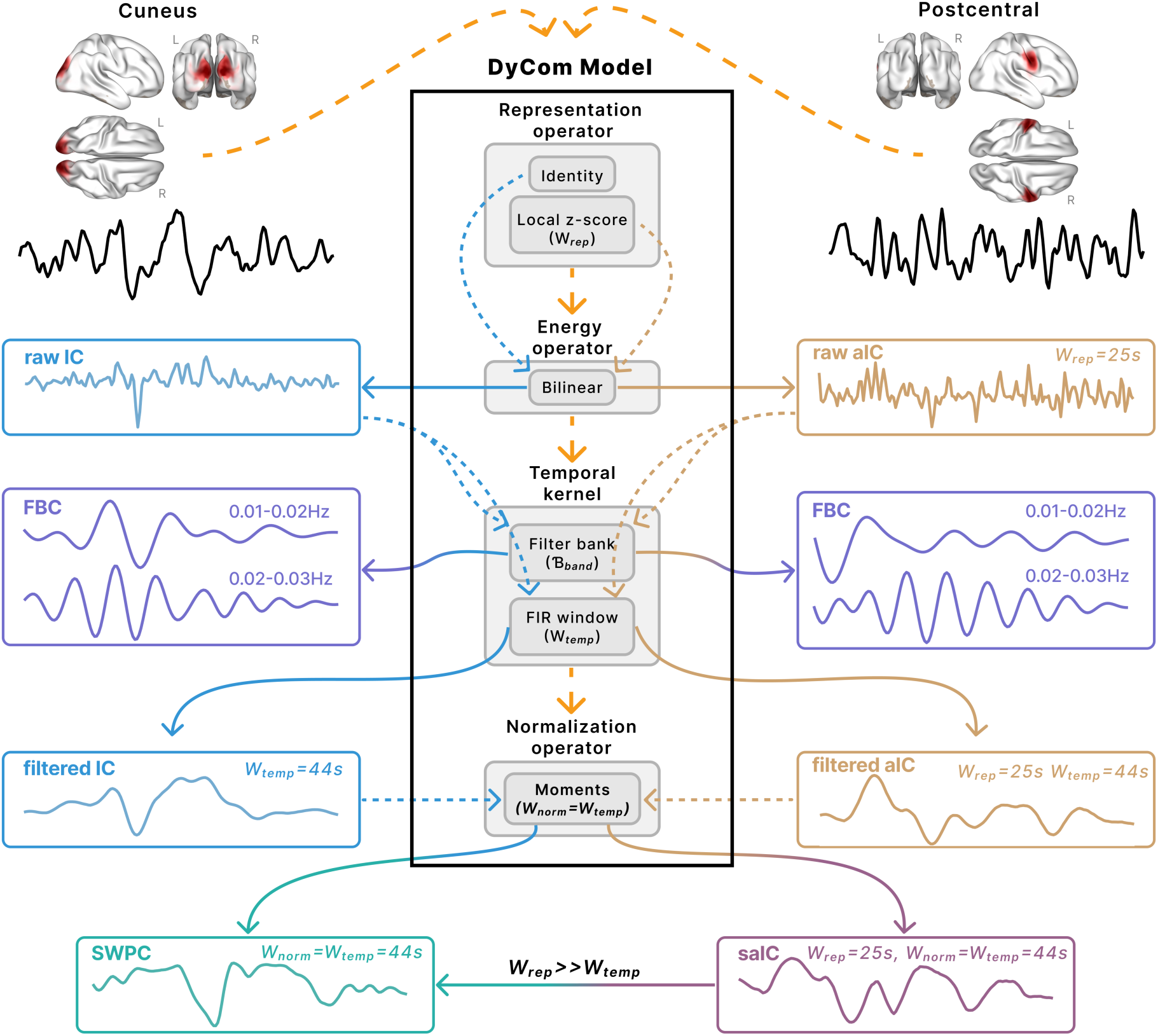
Modular structure of the DyCoM framework and its parameterizations. The DyCoM framework expresses dynamic connectivity as an ordered composition of four modular operators: representation, instantaneous energy, temporal integration, and normalization. The central schematic shows the signal processing flow within DyCoM, with solid arrows indicating how different estimators correspond to specific operator choices. Examples include instantaneous correlation (IC), sliding window Pearson correlation (SWPC), adaptive instantaneous correlation (aIC), standardized adaptive instantaneous correlation (saIC), and filter-bank connectivity (FBC). Each estimator reflects a unique configuration of operators, such as identity versus local z-scoring at the representation stage, FIR versus filter-bank temporal kernels, and the presence or absence of normalization. Time series panels illustrate the output of each method, with timescales labeled (e.g., W_temp = 44 s). This operator-level structure highlights the shared logic across methods and provides a common framework for comparison, interpretation, and systematic development of new estimators.

#### 2.1.1. Representation

Let *x_i_*(*t*) and *x*_j_(*t*) denote two time series drawn from an ℳ-dimensional multivariate signal. Each signal is first transformed through a representational operator ℛ[·], which defines the representation to be analyzed:

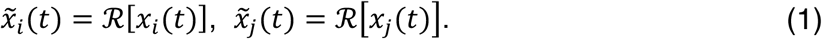

The representational operator **R**[·] is intentionally flexible and can take many forms depending on the analytical goal. It may act as an identity transform, apply local or adaptive normalization, extract analytic or amplitude-envelope representations, emphasize nonlinear energy terms, or isolate specific frequency components. By selecting **R**[·], DyCoM defines the feature space in which interactions are quantified. Representative forms of the representational operator are summarized in Table 1. Each choice of **R**[·] specifies the signal representation that contributes to the instantaneous energy operator introduced in the next stage.

**Table 1.**
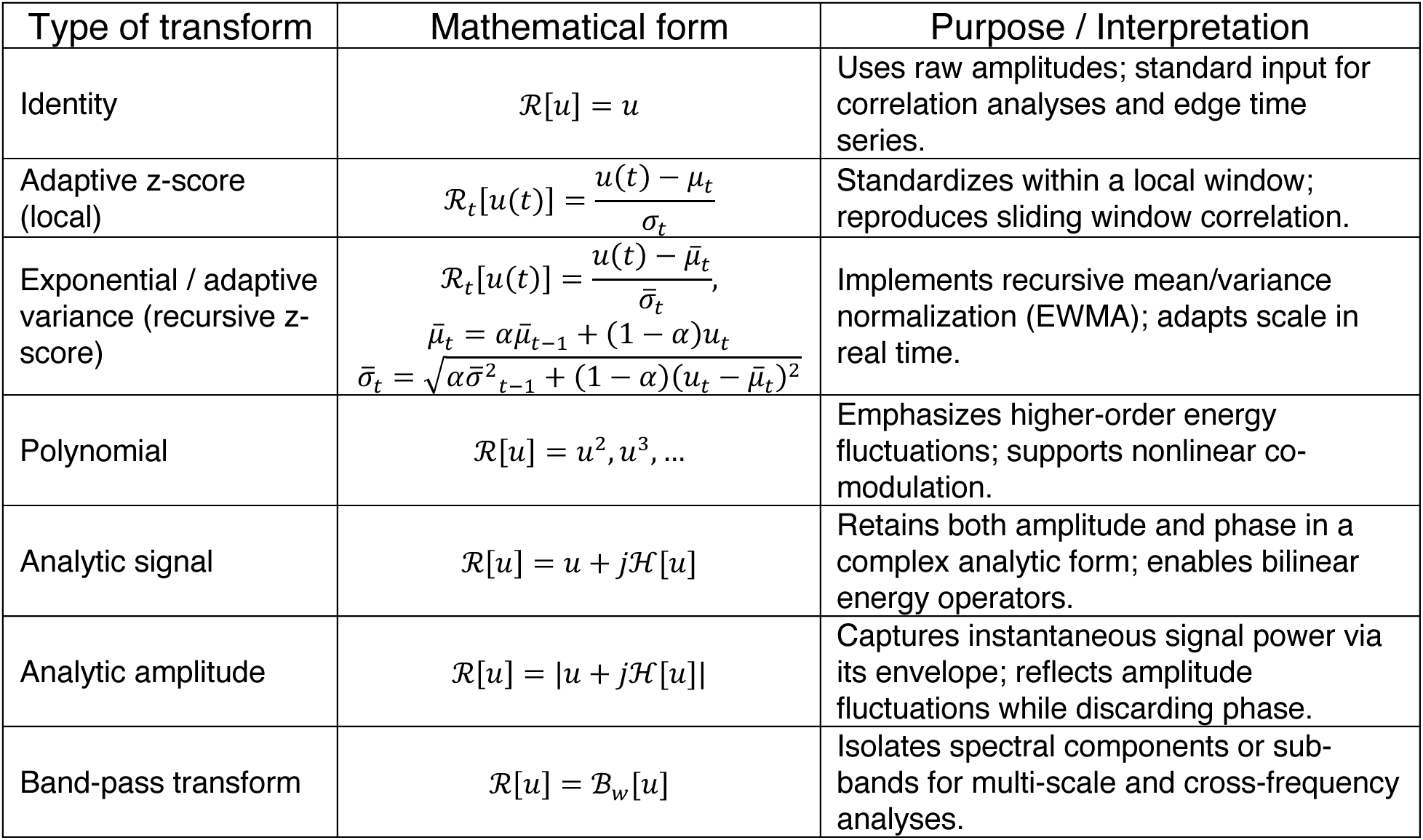
Representative forms of the representational operator R[·].

#### 2.1.2. Instantaneous energy kernel

At each time *t*, the instantaneous interaction between two representational signals is captured by an energy operator that defines how their instantaneous magnitudes or complex analytic representations combine to produce a local measure of co-modulation energy:

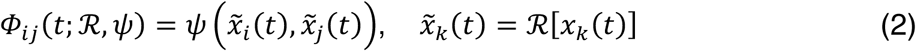

Here, the mapping *ψ*(·,·) specifies how two signals transformed by the representation operator interact at a single moment. Different choices of *ψ* describe different forms of instantaneous coupling, ranging from linear products to nonlinear or similarity-based measures.

A particularly important case is the bilinear energy operator,

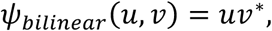

which yields the most fundamental bilinear interaction between two signals and quantifies their instantaneous co-modulation energy. In classical signal processing, this term corresponds to the zero-lag component of the cross Wigner-Ville distribution and represents the time-domain marginal of a bilinear energy operator (see Supplementary)^23^.

For representational operators that yield real-valued signals, the conjugate term reduces to *uv*^∗^ = *uv*, recovering the instantaneous correlation signal commonly used in dynamic functional connectivity. For analytic or complex representations, the Hermitian form *x^~^_i_(t)*. *x^~*^_j_(t)* preserves both amplitude and phase information, maintaining compatibility with established bilinear time-frequency energy formulations. The bilinear case, therefore, provides a conceptually grounded and computationally simple link between DyCoM and classical energy-based signal analysis. Beyond the bilinear case, nonlinear choices of *ψ* extend instantaneous co-modulation to higher-order or non-Euclidean relationships while preserving the same operator structure (Table 2).

**Table 2.**
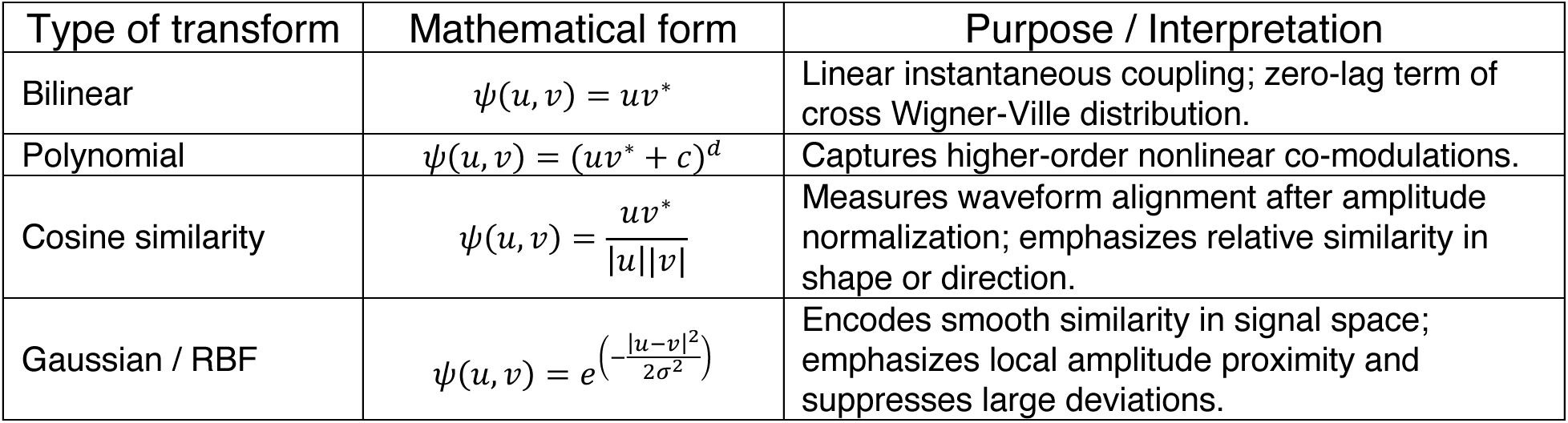
Representative forms of the instantaneous energy operator *ψ*(·,·)

**Table 3.** Representative forms of the temporal kernel ℎ(*τ*)

| Type of transform | Mathematical form | Purpose / Interpretation |
| --- | --- | --- |
| Identity | $h[\tau] = \delta[\tau]$ | Purely instantaneous co-modulation; equivalent to instantaneous correlation, edge time series or zero-memory case. |
| Rectangular / tapered (FIR) | $h[\tau] = \frac{\omega[\tau]}{\sum \omega[\tau]}, \omega[\tau] \in$<br>can be Rectangular, Hann, Hamming, Blackman, etc. | Fixed length averaging with optional spectral tapering; controls edge effects and frequency leakage, equivalent to an SWPC analysis with different window types. |
| Gaussian (FIR / Continuous) | $h[\tau] = e^{-\frac{\tau^2}{2\sigma^2}}$ | Smooth, symmetric weighting with minimal spectral leakage; acts as a probabilistic or diffusion-like temporal smoother, equivalent to an SWPC with a Gaussian window. |
| Exponential (IIR) | $h[\tau] = \alpha(1 - \alpha)^\tau, \tau \geq 0$ | Adaptive integration with fading memory; enables recursive real-time updates and adaptive filtering. |
| Multi-scale / Filter-bank | $h_\ell[\tau] = \mathcal{F}^{-1}\{H_\ell(f)\}$ where $H_\ell(f)$ is the filter's frequency response | Captures co-modulation dynamics across multiple temporal scales for multi-resolution analysis. |

**Table 4.**
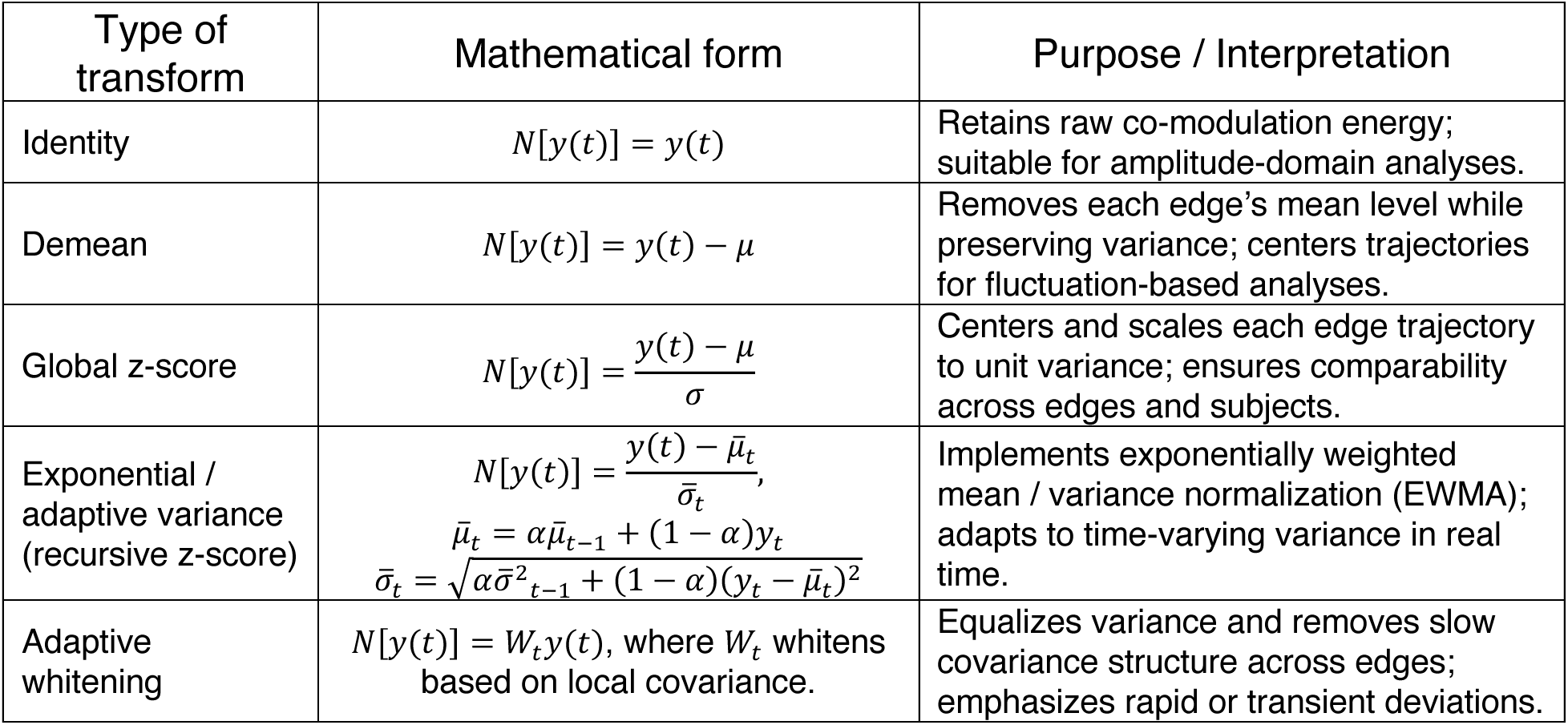
Representative forms of the normalized operator *N_t_*[⋅].

Each choice of *ψ* defines how two signals co-modulate at a single time point, providing the instantaneous energy structure from which temporal integration derives timescale-dependent dynamics.

#### 2.1.3. Timescale integration (filtering and adaptive weighting)

The instantaneous energy kernel Φ*_i_*_j_(*t*; ℛ, *ψ*) captures co-modulation at the finest temporal resolution. To characterize how these interactions evolve over time, DyCoM applies a temporal kernel ℎ(*τ*) that integrates instantaneous energy across a defined window or memory span:

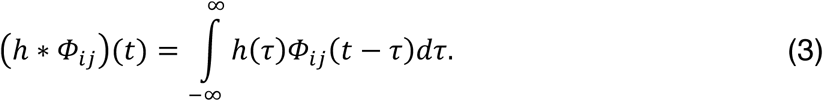

For linear representations and energy mappings, equivalent filtering could be applied directly to the original time series. However, applying ℎ(*τ*) after interaction formation provides greater flexibility and naturally extends to nonlinear energy operators. This formulation allows DyCoM to control the temporal resolution of the interaction itself, preserving rapid signal fluctuations while selectively smoothing their joint dynamics.

The kernel ℎ(*τ*) determines the effective temporal scale of co-modulation. Short kernels emphasize transient interactions, whereas longer kernels integrate sustained coupling, with finite impulse response (FIR) windows reproducing sliding window behavior as in SWPC. Infinite impulse response (IIR) kernels, such as exponential decays, yield adaptive trajectories with fading memory and support recursive updates. Families of spectral or multi-band kernels further decompose co-modulation into distinct frequency bands or timescales, yielding a multi-resolution representation of co-modulation dynamics.

#### 2.1.4. Normalization

After temporal integration, DyCoM can optionally apply an edge-domain normalization operator *N_t_*[⋅] to express co-modulation energy on a standardized scale. While many normalization procedures, such as local or adaptive z-scoring, can be embedded directly within the representational operator ℛ[·], this final normalization acts on the temporally integrated edge trajectories themselves. Applying normalization after integration ensures comparability across time, regions, and subjects when the integrated co-modulation trajectories (ℎ ∗ Φ*_i_*_j_)(*t*) differ in magnitude or variance.

This step compensates for differences in co-modulation energy arising from unequal signal power, slow drifts, or nonstationary scaling, producing trajectories that are easier to compare or combine statistically. Different choices of *N_t_*[⋅] therefore yield distinct families of DyCoM measures. The identity operator leaves integrated energy unscaled; global or adaptive z-scoring produces standardized, correlation-like trajectories; and adaptive whitening removes slow covariance structure across edges to emphasize transient deviations. Together, these options align DyCoM outputs with different analytical or physiological goals while preserving a unified mathematical structure.

#### 2.1.5. Unified DyCoM operator and matrix formulation

Given the four components described above, dynamic co-modulation can be expressed as a unified operator acting on a pair of signals:

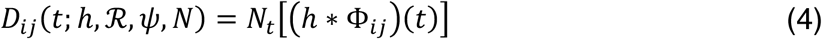

where

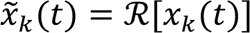 defines the representational signals,

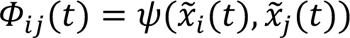 defines their instantaneous energy, and ℎ(*τ*) integrates energy across time to reveal the temporal structure of co-modulation. The normalization *N_t_*[⋅] is optional, used primarily to harmonize scales or accentuate transient deviations in the resulting co-modulation trajectories.

For multivariate data **x**(*t*) ∈ *F*^ℳ^, where F ∈ [ℝ, ℂ] denotes the underlying field (real or complex, respectively) depending on the chosen representation, the dynamic co-modulation process can be expressed compactly in matrix form. At each time point *t*, the representational feature vector is given by 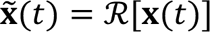, and the instantaneous interaction structure across all signal pairs is captured by the co-modulation kernel matrix

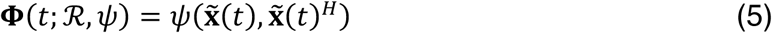

where *H* denotes the Hermitian (complex-conjugate transpose), which reduces to a standard transpose for real-valued data, and *ψ*(⋅,⋅) acts elementwise to generate pairwise energy interactions. In the bilinear case, this reduces to the outer product 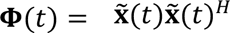, which defines the instantaneous Gram matrix of co-modulation energy.

Temporal integration is then applied to the full matrix trajectory to accumulate interaction information over the timescale defined by ℎ(*τ*), yielding (ℎ ∗ **Φ**)(*t*). Finally, a normalization operator *N_t_*[⋅] acts on the integrated matrix to ensure scale consistency across regions and time, producing the complete dynamic co-modulation representation

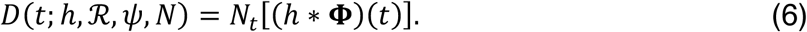

Each element *D_i_*_j_(*t*) reflects the normalized, timescale-specific co-modulation between signals *i* and *j*, while the full matrix *D*(*t*) forms a time-varying trajectory of network interactions. Together, Equations (1) – (6) formalize DyCoM as a unified operator family encompassing both instantaneous and temporally integrated measures of co-modulation, bridging classical energy-operator theory with contemporary dynamic connectivity analysis.

### 2.2. Relation to existing dynamic connectivity estimators

The DyCoM framework provides a unifying operator view of dynamic connectivity, showing that many previously independent estimators arise as special cases of a common underlying process. From this perspective, classical dynamic connectivity methods differ primarily in how the representation, energy mapping, temporal integration, and normalization operators are parameterized, rather than in their fundamental computational structure.

Within the correlation-based family of DyCoM parameterizations, several widely used methods, including instantaneous correlation, sliding window Pearson correlation, and filter-bank connectivity, share the same bilinear instantaneous energy operator but differ in how temporal integration and normalization are applied. Instantaneous correlation emerges as the simplest DyCoM case, with identity representation, bilinear energy mapping, and identity temporal integration and normalization^4^. The widely used sliding window Pearson correlation^1,3^ results when a finite impulse response temporal kernel is applied to the instantaneous products and to the local signal to generate moments for the moment-based normalization used in conventional correlation analyses. Filter-bank connectivity^5^ extends this structure by introducing multiple band-limited temporal kernels applied to the instantaneous interaction trajectories, enabling multiscale decomposition of co-modulation dynamics. In its original formulation, this approach implicitly applies local adaptive normalization to the signals prior to interaction formation^5^, but this operation is not explicitly named or treated as a standalone estimator. Under the DyCoM framework, this intermediate operation is revealed as a distinct representation-level transformation that precedes temporal integration.

In this work, we formalize this representation-level operation as a new estimator termed adaptive instantaneous correlation (aIC). The aIC applies local adaptive z-scoring to each signal prior to interaction formation, suppressing slowly varying or global components before the bilinear energy is computed. This formulation isolates moment-to-moment co-fluctuations while reducing shared drifts and physiological or vascular confounds (e.g., residual *CO*_2_ or respiration effects) that can persist after standard preprocessing of fMRI data^26,27^. By explicitly defining aIC within the DyCoM framework sequence, we elevate what was previously an implicit processing step into a principled, standalone dynamic connectivity measure.

We further define a recursive instantaneous correlation (rIC), which replaces local windowed normalization with exponentially weighted adaptive normalization, yielding an online-ready formulation that updates continuously in real time. rIC serves here to illustrate the framework’s extensibility to causal, real-time estimation; the analyses that follow focus on the offline parameterizations. Both aIC and rIC arise naturally from the DyCoM framework by modifying only the representation operator, while leaving the instantaneous energy mapping unchanged. Amplitude envelope correlation instantiates DyCoM using analytic amplitude representations^28,29^, while more recent nonlinear or kernelized approaches can also be expressed within the same operator framework by adopting alternative instantaneous energy mappings, such as polynomial, cosine, or Gaussian similarity functions^30,31^.

Representative dynamic connectivity estimators and their corresponding DyCoM parameterizations are summarized in Table 5. A schematic overview of these operator relationships is shown in Figure 1, which illustrates the DyCoM framework at the operator level and demonstrates how different parameterizations recover established methods while also generating new estimators. The figure is intended to emphasize the shared computational structure across methods rather than to highlight any single instantiation.

**Table 5.**
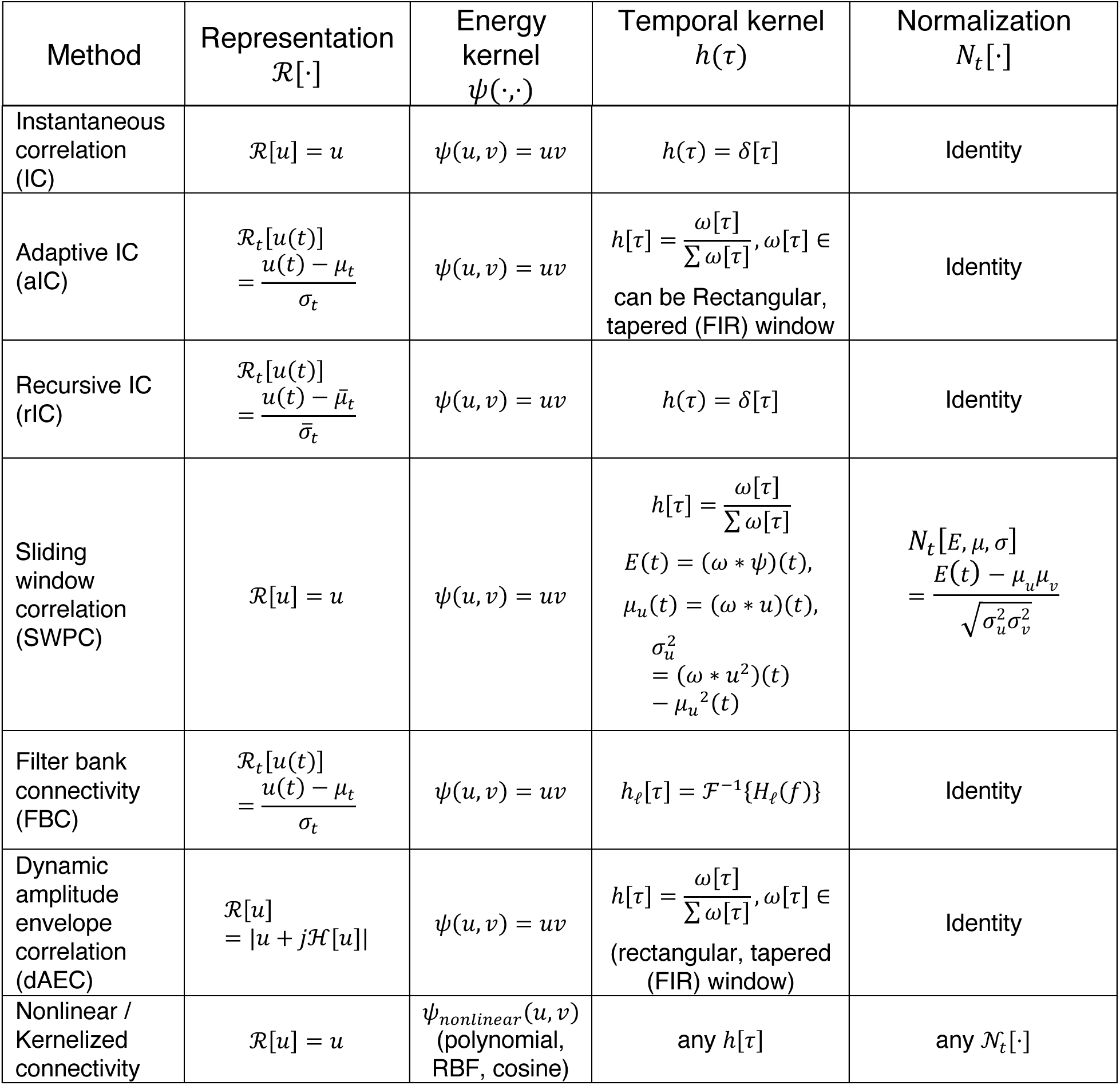
Representative dFNC methods as DyCoM parameterizations.

These mappings show that a broad class of amplitude-domain dynamic connectivity measures, ranging from instantaneous edge co-fluctuations to windowed and multiscale correlations, share a common mathematical identity under the DyCoM framework. Crucially, DyCoM not only unifies existing approaches but also exposes previously implicit operations and enables the principled construction of new dynamic connectivity estimators.

### 2.3. Standardized Adaptive Instantaneous Correlation (saIC)

Within the correlation family of DyCoM parameterizations, different estimators arise primarily from how representation and normalization are applied within the operator chain. Existing approaches do not combine adaptive local standardization at the representation stage with moment-based normalization at the interaction level. As a result, current methods either remove time-varying nuisance effects without enforcing bounded, interpretable scaling, or enforce correlation-style normalization without adaptively suppressing shared low frequency variance before interaction formation.

To address this gap, we introduce the standardized adaptive instantaneous correlation (saIC), a DyCoM parameterization that instantiates all four operators of the framework in non-identity form. The saIC is thus a principled combination of operations that exist individually in prior pipelines; the framework’s contribution is to make this combination explicit, bounded, and analyzable. The saIC method applies adaptive local z-scoring at the representation stage to remove shared global variance and slowly varying components before the bilinear interaction is formed. After temporal integration, it applies moment-based normalization at the interaction level, yielding trajectories that are expressed on a standardized correlation scale bounded between −1 and 1.

Formally, for saIC the DyCoM operators in (6) take the following forms:

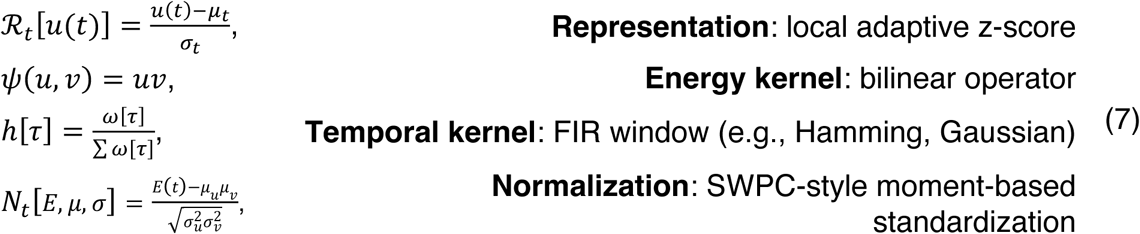

This configuration preserves the physiological interpretability of adaptive preprocessing while maintaining the statistical comparability and boundedness of correlation-based measures. As illustrated in Figure 1, saIC provides a concrete realization of the full DyCoM operator pipeline while remaining directly connected to established estimators through simple operator choices. Specifically, setting individual operators to the identity recovers commonly used methods as special cases. Setting the normalization operator to identity recovers aIC, which can then be combined with multi-band temporal kernels to produce filter-bank connectivity. Disabling the representational layer yields SWPC, and disabling both representation and normalization yields IC, with optional temporal filtering producing filtered IC variants. In this sense, saIC functions as a superset formulation from which classical correlation-based dynamic connectivity estimators emerge through principled operator simplifications.

Importantly, saIC also subsumes SWPC as a limiting case. Because the normalization layer is always defined by moments computed over the temporal kernel, when the representational window is larger than the temporal kernel window, the representational mean and variance change slowly relative to the normalization window and are therefore effectively constant within it. In this regime, saIC approaches the classical SWPC formulation without disabling any operators. This equivalence is demonstrated analytically in the Supplementary Material and illustrated in Figure 1.

The emergence of saIC highlights the generative nature of the DyCoM framework. By separating representation, interaction formation, temporal integration, and normalization into independently configurable operators, DyCoM not only recovers established estimators but also enables the principled construction of new, physiologically grounded dynamic connectivity measures that unify and extend existing approaches.

### 2.4. Simulations of DyCoM methods

To evaluate DyCoM parameterizations under controlled conditions with known ground truth, we constructed a set of simulations in which time-varying connectivity between two neural sources was explicitly prescribed. All simulation scenarios shared the same underlying neural coupling trajectory and differed only in the presence and structure of nuisance components, enabling direct comparison of estimator sensitivity to physiologically realistic confounds.

Two regional time courses, *x*(*t*) and *y*(*t*), were simulated at a sampling rate of *f_s_* = 1 Hz for a total duration of *T* = 2000 s. Their instantaneous correlation was defined by a single ground-truth connectivity trajectory *ρ*(*t*), generated by band-limiting a Gaussian white noise sequence ∼N(0,1) to the frequency range [0.002 0.004] Hz and scaling its amplitude to a maximum of 0.9. This strong, slowly varying trajectory reflects minute-scale fluctuations in resting-state functional coupling and corresponds to the regime in which correlation-based dynamic connectivity estimators perform optimally^17^.

At each time point, the underlying neural signals were generated such that their instantaneous covariance matched *ρ*(*t*). Two independent white-noise sources *u*_1_(*t*), *u*_2_(*t*) ∼ N(0,1) were first filtered to the canonical BOLD band (0.01–0.15 Hz). Their samples were then linearly mixed through a time-dependent covariance matrix

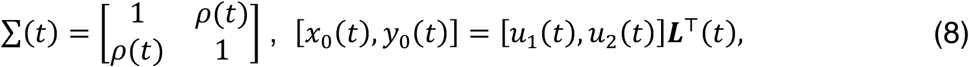

where ***L***(*t*) is the Cholesky factor of ***Σ***(*t*). This procedure yields two signals that precisely match the target correlation *ρ*(*t*) while preserving realistic BOLD frequency band structure.

Using this base neural signal model, we examined six simulation scenarios:

#### (i) Clean neural coupling

The observed signals were set equal to the neural components,

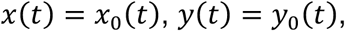

providing an idealized reference condition without nuisance contamination.

#### (ii) Weak shared nuisance

To emulate mild physiological confounds, we added a shared global signal *g*(*t*) generated as band-limited Gaussian noise in the low frequency range (0.02–0.04 Hz), mimicking respiration- or CO₂-driven vasodilatory oscillations. The amplitude of the global component was scaled such that approximately 20% (*λ* = 0.20) of each node’s variance originated from the shared signal.

#### (iii) Strong shared nuisance

This condition was identical to the weak shared nuisance case, except that the global component accounted for approximately 60% (*λ* = 0.60) of each node’s variance, modeling severe systemic contamination.

#### (iv) Strong unshared nuisance

To emulate the presence of multiple independent or region-specific confounds, we added strong nuisance components that were not fully shared between regions. This scenario reflects situations in which motion, vascular, or scanner-related artifacts project differently across brain regions, despite standard preprocessing.

#### (v) Slow neural coupling with region-specific nuisance

To examine the regime in which the coupled neural fluctuations themselves occupy low frequencies, the neural sources were band-limited to 0.01–0.08 Hz, placing most of the coupled signal content below the adaptive representational cutoff, while contamination was restricted to fast, region-specific physiological components. Independent nuisance signals, band-limited to 0.09–0.11 Hz and generated separately for each node, were scaled such that approximately 25% (λ = 0.25) of each node’s variance originated from nuisance, with a small region-specific asymmetry as in the shared-nuisance scenarios.

#### (vi) Time-varying connectivity at nuisance timescales

To create genuine spectral ambiguity between coupling dynamics and contamination, the ground-truth trajectory ρ(t) was band-limited to 0.01–0.03 Hz, directly overlapping the frequency range of the shared nuisance (0.02–0.04 Hz), which was applied at the strong level (λ = 0.60) as in Scenario 3. In this regime, nuisance-driven covariance fluctuations occupy the same timescales as the prescribed connectivity dynamics and cannot be separated by any spectral margin.

For scenarios involving shared nuisance components, the global signal was scaled according to a variance-fraction rule,

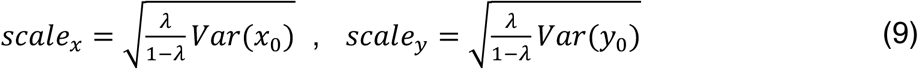

and a small asymmetry was introduced to reflect region-specific susceptibility,

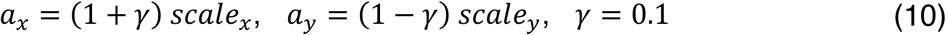

The observed signals were therefore given by:

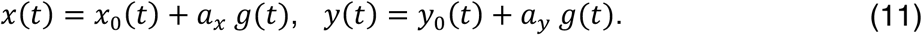

Before analysis, all time series were z-scored to remove mean offsets and ensure consistent scaling across methods.

Each signal pair was analyzed using four DyCoM parameterizations: instantaneous correlation, sliding window Pearson correlation, adaptive instantaneous correlation, and standardized adaptive instantaneous correlation. All DyCoM parameterizations in this study employed a bilinear energy operator *ψ*(*u*, *v*) = *uv* and a Hamming FIR temporal kernel of 88 s for temporal integration. The kernel was set to correspond to the −3 dB cutoff at 0.01 Hz, reflecting the lowest frequency retained in the simulated data and matching the lowest reliably observable frequency in the resting-state BOLD signal^5,17,25^. Methods incorporating adaptive representation (aIC and saIC) additionally required a representational-layer window for local z-scoring. This window determines the lowest-frequency nuisance components that can be effectively suppressed and was set to 0.03 Hz, corresponding to a window length of 33 s, so that its cutoff lies within the simulated nuisance band (0.02–0.04 Hz) and suppresses the slower portion of the shared nuisance (see Supplementary Fig. S2 for sensitivity of the results to both window choices). Two scenario-specific exceptions were made to match the estimators to the prescribed dynamics. In Scenario 5, the representational window was set to 17 s (cutoff ≈ 0.06 Hz), placing the adaptive cutoff above the coupled neural band, and in Scenario 6 the temporal kernel was shortened to 22 s so that the integration timescale could track the faster connectivity trajectory, with all other operator settings unchanged.

Each simulation scenario was repeated for 1000 independent trials using different random seeds while preserving the same ground-truth trajectory *ρ*(*t*), Estimated connectivity trajectories *ρ^^^(t)* were compared to the ground truth using Pearson correlation and root-mean-square error. Mean estimates and 95% pointwise confidence intervals were computed across trials to quantify estimator accuracy and variability under each nuisance regime.

### 2.5. fMRI data

Resting-state fMRI data were obtained from the Function Biomedical Informatics Research Network (fBIRN), comprising individuals with schizophrenia (SZ) and matched healthy control subjects (CN). Imaging data were acquired with a repetition time (TR) of 2 s. The final sample included 160 CNs (mean age 37.04 ± 10.86 years, range 19–59; 45 female) and 151 SZ patients (mean age 38.77 ± 11.63 years, range 18–62; 36 female). The two groups were well matched with respect to age (p = 0.18), sex distribution (p = 0.39), and mean framewise displacement during scanning (p = 0.97). All SZ participants were clinically stable at the time of data acquisition. This dataset was selected to evaluate the clinical sensitivity and interpretability of DyCoM-based dynamic connectivity estimators in a well-characterized psychiatric population. Data collection was approved by the institutional review boards of the participating fBIRN sites, with written informed consent obtained from all participants; the present study analyzed the data in de-identified form.

### 2.6. fMRI data processing

Resting-state fMRI data underwent standard preprocessing to mitigate noise and imaging artifacts prior to analysis. Preprocessing was performed using Statistical Parametric Mapping (SPM5; http://www.fil.ion.ucl.ac.uk/spm)^32^ and included slice-timing correction, rigid-body realignment, spatial normalization to standard space, and spatial smoothing, following established pipelines^33^. Following preprocessing, subject-level intrinsic connectivity networks (ICNs) were extracted using the NeuroMark framework, a fully automated spatially constrained independent component analysis (ICA) approach^34^. NeuroMark leverages population priors to ensure robust and reproducible network estimation across subjects. Specifically, we employed the neuromark_fMRI_1.0 template (available within the GIFT toolbox at http://trendscenter.org/software/gift), yielding 53 ICNs per subject. These components span seven functional domains: Subcortical (SC), Auditory (AUD), Sensorimotor (SM), Visual (VIS), Cognitive Control (CC), Default Mode (DM), and Cerebellar (CB).

To further enhance signal quality, ICN time courses were detrended and despiked to remove residual low frequency drifts, abrupt signal excursions, and motion-related artifacts that may persist after preprocessing. The ICN time series were then bandpass filtered in the range 0.01–0.15 Hz, a frequency band widely regarded as relevant for capturing physiologically meaningful BOLD fluctuations in resting-state fMRI^35,36^. Filtering was implemented using a zero-phase infinite impulse response Butterworth filter designed with MATLAB’s *butter* function and applied using *filtfilt* to avoid phase distortion^37^. The filter order was selected using the *buttord* function tuned to maintain passband ripple below 3 dB and ensure at least 30 dB stopband attenuation. Finally, all ICN time series were z-scored to ensure consistent scaling across subjects and components. All analyses were performed in MATLAB R2025a.

### 2.7. Application of DyCoM to fMRI data

DyCoM analysis was applied to resting-state fMRI data using four parameterizations: instantaneous correlation, sliding window Pearson correlation, adaptive instantaneous correlation, and standardized adaptive instantaneous correlation. All methods were implemented within the DyCoM framework to ensure consistent operator definitions across analyses. To improve comparability across methods and to focus on differences arising from representation and normalization rather than timescale, all four estimators were evaluated using an explicit temporal integration stage, including IC and aIC, which do not require temporal smoothing in their canonical forms. Temporal integration was performed using a Hamming FIR window of 44 s, a window length widely used in prior fMRI studies to balance temporal resolution and statistical reliability^3,38^.

For methods incorporating adaptive representation (aIC and saIC), a representational-layer window was used to compute local mean and variance for adaptive z-scoring. This window was set to 33 s, corresponding to a cutoff frequency of approximately 0.03 Hz. This choice was motivated by prior work showing that BOLD signal components below approximately 0.03 Hz are strongly influenced by physiological and scanner-related confounds^39^, including CO₂-driven vasodilatory fluctuations and low frequency drift^40^, and by the widespread use of this cutoff as a boundary for narrowband conditions such as phase-based connectivity analyses in fMRI^7,17,41–43^. Setting the representational window to this timescale suppresses slowly varying nuisance components and reduces shared variance before interaction formation.

### 2.8. Temporal Integration flexibility within DyCoM

To illustrate the flexibility of the DyCoM framework with respect to temporal integration, we conducted a controlled sensitivity analysis in which only the temporal kernel was varied while all other operators were held fixed. Specifically, we compared three configurations: instantaneous correlation, 44 s finite impulse response integration, and 88 s finite impulse response integration. The 44 s window was selected to match the primary analysis and reflects a commonly used compromise between temporal resolution and estimation stability in resting-state fMRI. The 88s window was included as a longer integration condition corresponding approximately to the −3 dB cutoff associated with a 0.01 Hz lower frequency bound, which is widely used as the high-pass limit in resting-state BOLD preprocessing^5,44^. This choice allows evaluation of state structure at a timescale aligned with the slowest reliably retained physiological fluctuations in the data. In each case, the representation operator, instantaneous energy mapping, and normalization procedure were identical, ensuring that any differences in estimated states arose solely from the timescale of integration

For each temporal configuration, DyCoM edge trajectories were computed for all subjects and analyzed using the same preprocessing, dimensionality reduction, and clustering procedures described above. State centroids were estimated independently for each integration scale. To ensure consistent labeling across scales, states were matched based on the maximum Pearson correlation between centroids. To quantify domain-level organization, we computed a domain segregation contrast for each state and scale, defined as the difference between the mean absolute within-domain and between-domain connectivity magnitudes divided by their sum, with self-connections excluded. This procedure enabled assessment of how the temporal integration scale modulates the spatial configuration and domain structure of DyCoM-derived states while preserving a consistent operator formulation.

### 2.9. Dynamic analysis

For each DyCoM parameterization (IC, SWPC, aIC, and saIC), subject-specific dynamic connectivity trajectories were first demeaned to remove the static component and emphasize temporal fluctuations^45^. This centering was performed independently for each subject and method to isolate dynamic variability without confounding from mean connectivity differences. Dimensionality reduction was then applied separately for each method using principal component analysis. The number of retained components was selected using the elbow criterion. The retained components were clustered using k-means, with the number of clusters varied from 2 to 10. K-means was run with 20 random initializations and a maximum of 10,000 iterations to ensure stable convergence. For all methods, the optimal number of clusters was determined to be five based on the elbow criterion. This selection was performed separately for each parameterization, and the optimum was identical across all four (Supplementary Fig. S3).

Using the resulting state sequences, three standard dynamic connectivity metrics were computed for each participant. Mean dwell time (MDT) quantified state persistence as the average duration spent in a state once entered^46^. The fraction rate (FR) quantified state occupancy as the proportion of the total scan duration assigned to each state^46^. State-to-state transition probabilities were summarized in a transition matrix (TM), capturing the likelihood of transitioning between pairs of states^46^.

Group differences between healthy controls and individuals with schizophrenia, as well as associations with cognitive performance measured by the Computerized Multiphasic Interactive Neurocognitive System (CMINDS) and symptom severity measured by the Positive and Negative Syndrome Scale (PANSS), were assessed using generalized linear models. All models included age, sex, scanning site, and mean framewise displacement as covariates. Multiple comparisons were controlled using the Benjamini–Hochberg false discovery rate procedure. Correction was applied separately for each parameterization and each summary metric, that is, across the five state-specific comparisons for mean dwell time and fraction rate, and across the state-pair transitions for the transition matrix. Covariate-adjusted effect sizes and partial R² values are reported throughout.

## 3. RESULTS

### 3.1. Controlled simulations with known ground truth

We evaluated DyCoM parameterizations under controlled conditions with known ground truth to assess how operator choices influence sensitivity to shared and unshared nuisance components. All methods were compared against the same prescribed time-varying connectivity trajectory, and performance was quantified across 1000 trials using correlation with the ground truth and root-mean-square error. Differences across scenarios, therefore, reflect estimator robustness rather than differences in underlying neural coupling.

Under clean conditions without nuisance signals (Scenario 1; Fig. 2a), all methods accurately recovered the ground-truth connectivity trajectory, exhibiting high correlation and low RMSE, establishing a common baseline of unbiased performance. As shared nuisance components were introduced, systematic differences emerged. With weak shared nuisance (Scenario 2; Fig. 2b), IC showed clear degradation, while SWPC provided partial robustness but remained influenced by a shared low-frequency variance. In contrast, aIC and saIC achieved the highest correspondence with the ground truth, yielding both the highest correlation and lowest RMSE. These trends became more pronounced under strong shared nuisance (Scenario 3; Fig. 2c), where IC showed substantial failure to recover the underlying trajectory, and SWPC showed only modest improvement, whereas aIC and saIC remained stable, with saIC consistently achieving lower RMSE.

**Figure 2:**
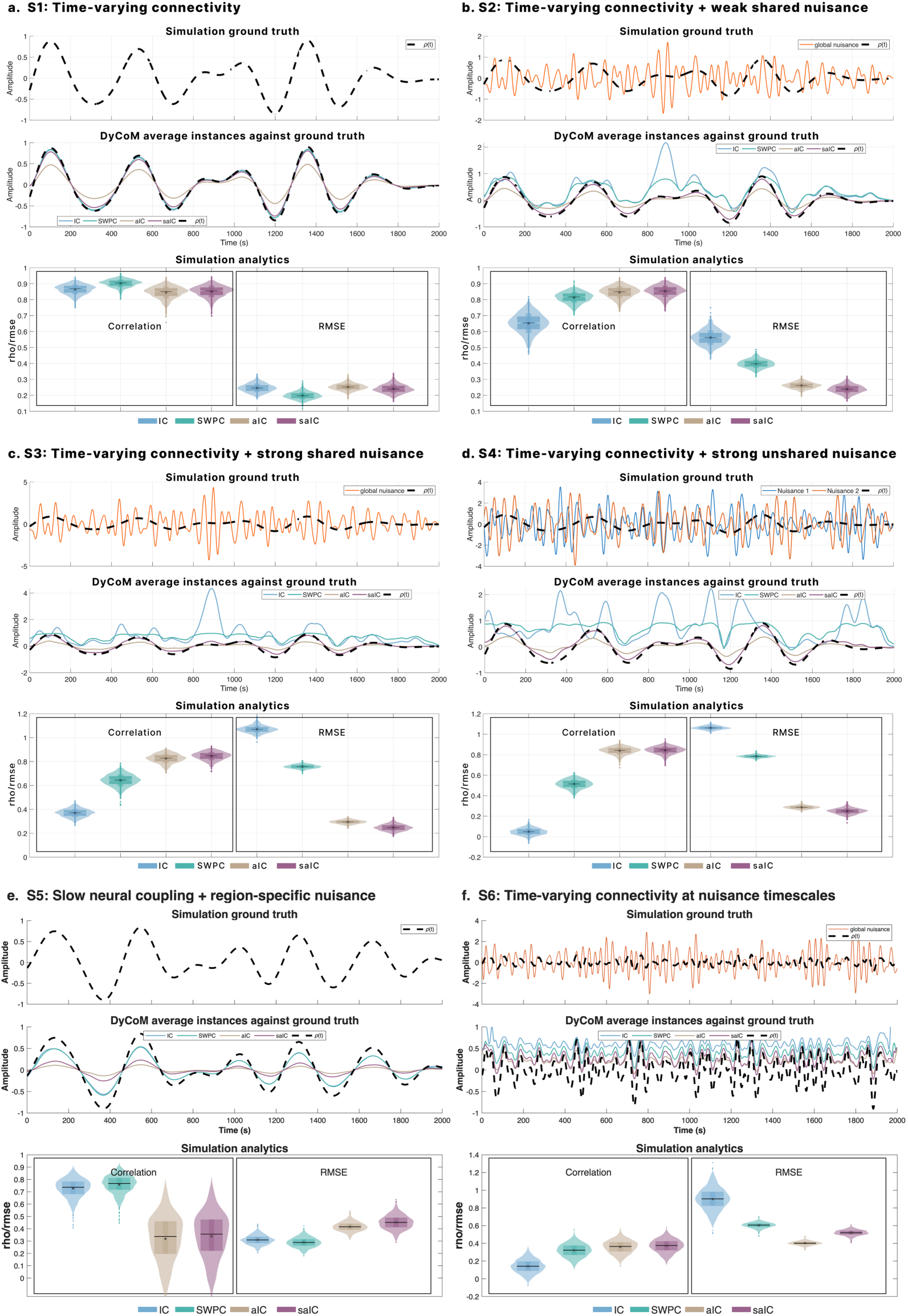
DyCoM simulations reveal estimator sensitivity to shared and unshared nuisance. Simulation scenarios evaluate four DyCoM parameterizations: IC, SWPC, aIC, and saIC, under different increasing and varying levels of nuisance. In each panel, the top row shows the ground truth connectivity trajectory (ρ(t)) with or without simulated noise. The middle row plots average estimated trajectories across 1000 trials, overlaid with the ground truth. The bottom row presents violin plots of estimator performance, quantified by Pearson correlation and root-mean-square error (RMSE) relative to ρ(t). **a.** Scenario 1 shows that all estimators perform similarly under clean conditions. **b.** In Scenario 2, weak shared nuisance degrades IC, while aIC and saIC remain robust. **c.** Scenario 3 introduces a strong shared nuisance, leading to further breakdown in IC and SWPC, while aIC and saIC maintain high accuracy. **d.** Scenario 4 simulates a strong unshared nuisance, where the aIC and saIC methods continue to perform well. **e.** Scenario 5 prescribes slow neural coupling with region-specific physiological nuisance, a regime in which IC and SWPC are the more accurate estimators, while the adaptive representation removes the slow fluctuations that carry the coupling. **f.** Scenario 6 places the connectivity dynamics at the same timescales as a strong shared nuisance, where spurious and true co-modulation become inseparable, and no estimator recovers the ground truth. These results demonstrate that estimator robustness depends on operator composition rather than estimator identity, with each parameterization possessing a distinct regime of validity.

When nuisance effects were strong and unshared across regions (Scenario 4; Fig. 2d), IC and SWPC again showed marked degradation, particularly in RMSE, reflecting sensitivity to asymmetric confounds. Both aIC and saIC remained comparatively invariant under this condition, with saIC retaining the lowest RMSE overall. In Scenario 5 (Fig. 2e), where the coupled neural fluctuations were slow and contamination was limited to fast, region-specific physiological nuisance, the pattern inverted: IC and SWPC recovered the trajectory most accurately, whereas aIC and saIC removed the slow fluctuations carrying the coupling and showed reduced recovery, identifying a regime in which the non-adaptive parameterizations are preferred. In Scenario 6 (Fig. 2f), the connectivity dynamics occupied the same timescales as a strong shared nuisance, rendering spurious and genuine co-modulation spectrally indistinguishable; no estimator recovered the ground truth reliably, delineating an identifiability boundary predicted by the framework, in which nuisance suppression requires a spectral margin between confound and coupling dynamics. Together with Scenarios 1–4, these results demonstrate the importance of the DyCoM operator layers in shaping robustness during dynamic connectivity estimation, with each parameterization possessing a distinct regime of validity. These conclusions were stable across window choices, additional nuisance regimes, and implementation checks (Supplementary Figs. S1–S2, Table S1).

### 3.2. Group differences in dynamic connectivity

We first assessed the sensitivity of DyCoM parameterizations to group differences between controls and individuals with schizophrenia using three dynamic metrics: mean dwell time, fraction rate, and transition matrix statistics. Figure 3a summarizes covariate-adjusted effect sizes (Cohen’s *D*), FDR-corrected significance levels, and partial R² values across methods.

**Figure 3:**
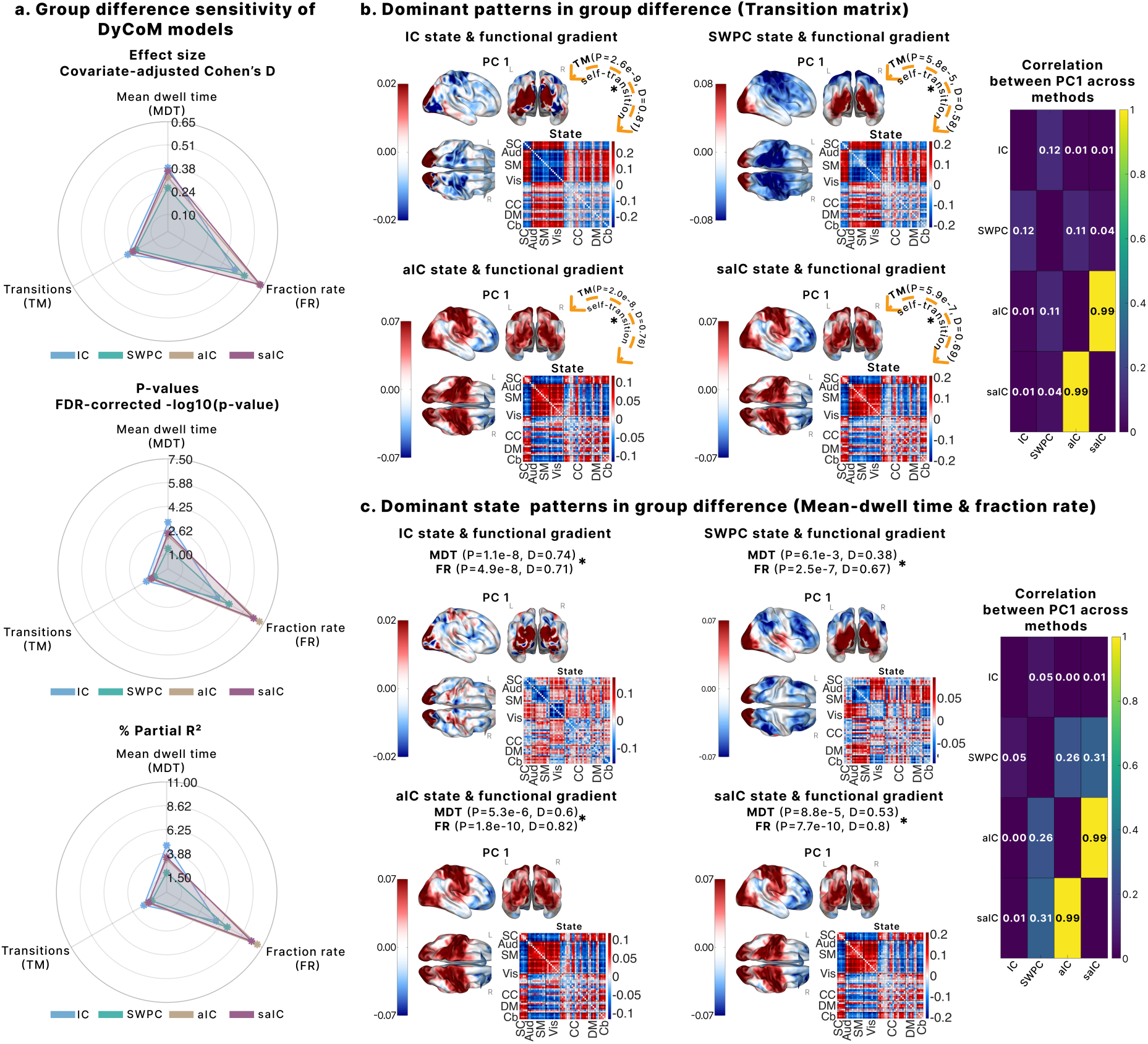
DyCoM parameterizations reveal complementary group differences in schizophrenia. **a.** Radar plots summarize effect sizes (covariate-adjusted Cohen’s D), statistical significance (FDR-corrected p-values), and partial R² for group differences in three dynamic metrics: mean dwell time (MDT), fraction rate (FR), and transition matrix statistics (TM). Each DyCoM parameterization shows distinct sensitivity across metrics. **b.** Dominant transition-related connectivity states from each method are visualized using their first principal component (PC1), overlaid on cortical surfaces and connectivity matrices. PC1 captures the spatial pattern driving group differences in TM. The rightmost matrix shows correlations between PC1 vectors across methods, quantifying similarity in dominant state structure. **c.** Dominant states for MDT and FR metrics are similarly projected onto cortical space, with PC1 again capturing the primary spatial mode of variation. The asterisks indicate an FDR corrected p-value<0.05. These results indicate that DyCoM parameterizations emphasize distinct neurobiological signatures of schizophrenia.

Distinct sensitivity profiles emerged across parameterizations and metrics. IC exhibited the largest group differentiation for MDT and TM, showing higher effect sizes, greater explained variance, and stronger statistical significance for these measures. In contrast, aIC and saIC showed the greatest sensitivity for FR, yielding the largest effect sizes and partial R² values for state occupancy. SWPC generally demonstrated reduced sensitivity across metrics relative to the other parameterizations. These results indicate that different DyCoM parameterizations emphasize complementary aspects of group-level dynamic differences rather than a single uniformly dominant estimator.

To characterize the network-level patterns underlying these group differences, we examined the connectivity states that drove sensitivity for each DyCoM parameterization. For each method, the state exhibiting the largest covariate-adjusted Cohen’s *D* across the dynamic metrics (MDT, FR, and TM) was selected as the dominant state. The spatial structure of each dominant state was summarized by its leading principal component (PC1), which captures the dominant node-wise expression of connectivity variation within that state. This component was projected onto cortical surfaces to visualize regional contributions to group differentiation.

Examination of the dominant transition-related states revealed method-specific but interpretable spatial organizations (Fig. 3b). Because dynamic trajectories were demeaned prior to clustering, states reflect fluctuations around a zero-centered baseline, such that consistent opposite polarities around the mean are identified as distinct dynamical states. IC exhibited a heterogeneous pattern in which higher-order visual regions, such as the cuneus and lingual cortex, loaded positively, while early visual cortex (calcarine), parietal, and sensorimotor regions loaded negatively. In contrast, SWPC produced a smooth posterior–anterior organization characterized by positive loadings in visual and temporal cortex and negative loadings across parietal, sensorimotor, and frontal regions. Both aIC and saIC converged on a highly coherent pattern, with strong positive loadings in visual, parietal, and sensorimotor regions and negative loadings in frontal and subcortical areas. The close correspondence between aIC and saIC reflects similar dominant connectivity patterns under adaptive parameterizations.

For mean dwell time and fraction rate, the dominant states identified for aIC and saIC closely matched those observed for transition probabilities, exhibiting nearly identical spatial organization (Fig. 3c). This consistency indicates that, for adaptive parameterizations, group differences across dynamic metrics are driven by differential expression of the same underlying connectivity state rather than by metric-specific state selection. In contrast, IC was characterized by a highly asymmetric pattern dominated by inferior parietal cortex, contrasted with positive loadings in visual regions, resulting in a fragmented spatial profile. SWPC produced a smoother posterior–anterior contrast, with positive contributions from visual and temporal cortex and negative contributions from parietal, sensorimotor, and frontal regions. Collectively, these findings illustrate how different DyCoM parameterizations highlight complementary aspects of dynamic connectivity structure at the network level.

### 3.3. Cognitive and clinical associations of dynamic connectivity

We next examined how DyCoM parameterizations differed in their associations with cognitive performance and clinical symptom severity. Associations were assessed using generalized linear models relating dynamic connectivity metrics to CMINDS cognitive scores and PANSS symptom measures, with covariate-adjusted partial *R*^2^ values used to quantify explained variance.

Figure 4a summarizes the cognitive association profiles across DyCoM parameterizations. Distinct patterns emerged across methods rather than uniform sensitivity. IC and SWPC showed their strongest associations with processing speed and the composite CMINDS score, with comparatively weaker relationships to learning and working memory domains. In contrast, aIC exhibited broader associations across verbal/visual learning, attention vigilance, and working memory, indicating sensitivity to higher-order cognitive functions. saIC showed more selective associations, with attenuated partial R² values across domains relative to aIC. These results indicate that DyCoM parameterizations emphasize different aspects of cognitive variability, with operator choices influencing which cognitive domains are most strongly captured, while also revealing partial overlap in association profiles across methods.

**Figure 4:**
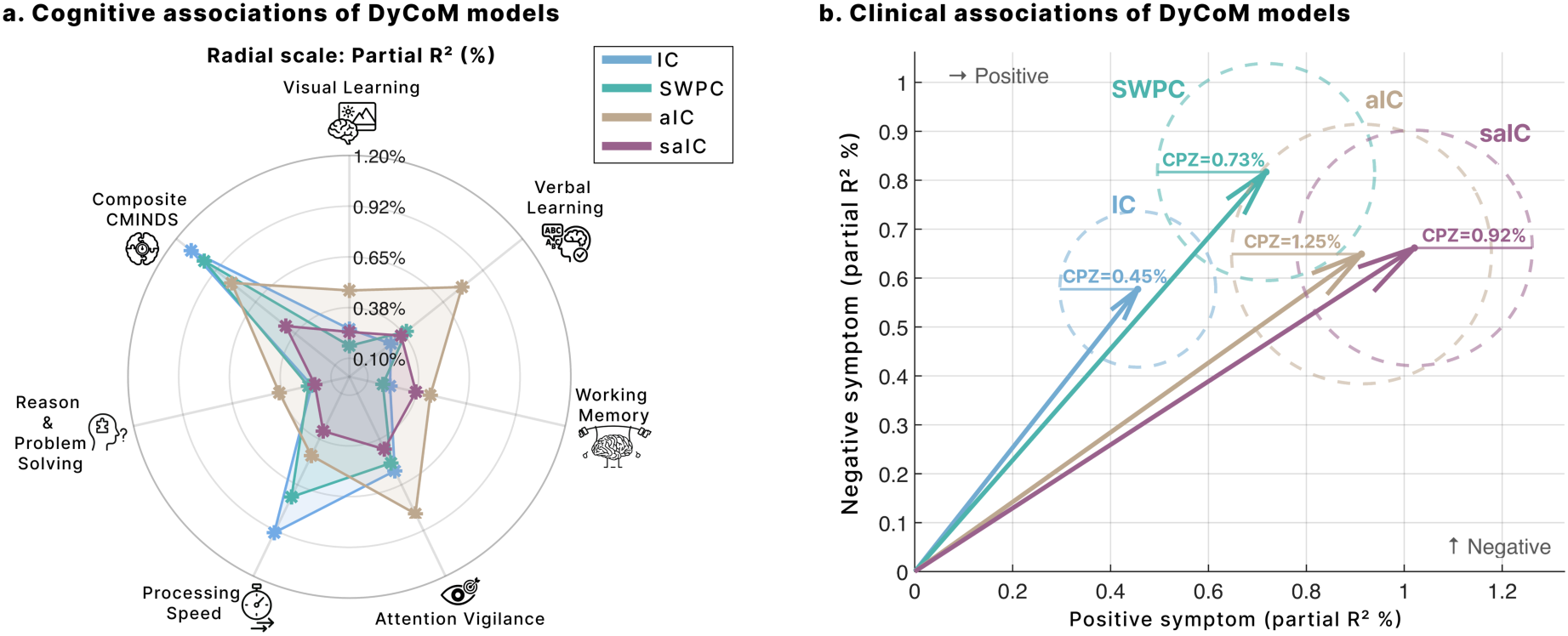
Cognitive and clinical associations of DyCoM parameterizations. **a.** Radar plots show partial R² values for associations between dynamic connectivity metrics and cognitive performance across seven domains from the CMINDS battery. IC and SWPC emphasize processing speed and composite scores, while aIC captures broader associations, particularly with learning and attention vigilance. **b.** Arrows indicate the relative strength of associations between dynamic connectivity and PANSS symptom dimensions (positive vs. negative), with arrowheads marking the vector tip and partial R² encoded by length. Colored dashed circles represent variance explained by antipsychotic medication dose (CPZ). IC and SWPC align more strongly with negative symptoms, while aIC and saIC exhibit more balanced associations and greater CPZ sensitivity. These profiles highlight how operator choices influence the clinical dimensions captured by each estimator.

Figure 4b illustrates associations with PANSS positive and negative symptom dimensions, with arrow direction indicating relative sensitivity and halo size indicating variance explained by antipsychotic dose equivalency (CPZ). The parameterizations occupied distinct regions in symptom-association space. IC and SWPC showed relatively stronger associations with negative symptoms compared to positive symptoms, accompanied by smaller CPZ-related variance. In contrast, aIC and saIC exhibited increased sensitivity to positive symptoms while retaining substantial associations with negative symptom severity, and showed larger CPZ-associated effects. Notably, the transition from IC and SWPC to adaptive parameterizations was accompanied by a systematic shift in the balance between positive and negative symptom associations, as well as increased coupling with medication dosage, rather than a uniform increase in explained variance. These findings demonstrate that operator-level design choices within DyCoM systematically govern which behavioral and clinical dimensions are most prominently captured, enabling principled interpretation of dynamic connectivity estimates across cognitive and symptom domains.

### 3.4. Temporal Integration Flexibility Reveals Scale-Dependent States

To demonstrate the flexibility of DyCoM with respect to temporal integration, we compared instantaneous interaction with 44s and 88s windowed formulations while holding all other operators fixed (Figure 5). States 1, 2, and 5 showed high cross-scale spatial similarity, indicating that their connectivity configurations are largely invariant to the temporal integration options chosen. In contrast, states 3 and 4 displayed marked scale dependence, with reduced similarity between instantaneous and windowed estimates, particularly at the longer 88 s integration (Fig. 5b). Domain-level organization reflected a similar pattern. For states 3 and 4, temporal integration increased within-domain dominance relative to between-domain connectivity, whereas states 1 and 2 remained near zero across scales. This pattern indicates that longer temporal integration accentuates coherent domain-level structure that is less pronounced in the instantaneous formulation. In contrast, instantaneous interaction emphasizes rapid co-fluctuations that are less organized into clear within-domain patterns. Together, these findings demonstrate that DyCoM flexibly spans instantaneous and windowed regimes within a unified operator structure, while revealing that certain connectivity states exhibit intrinsic scale dependence.

**Figure 5:**
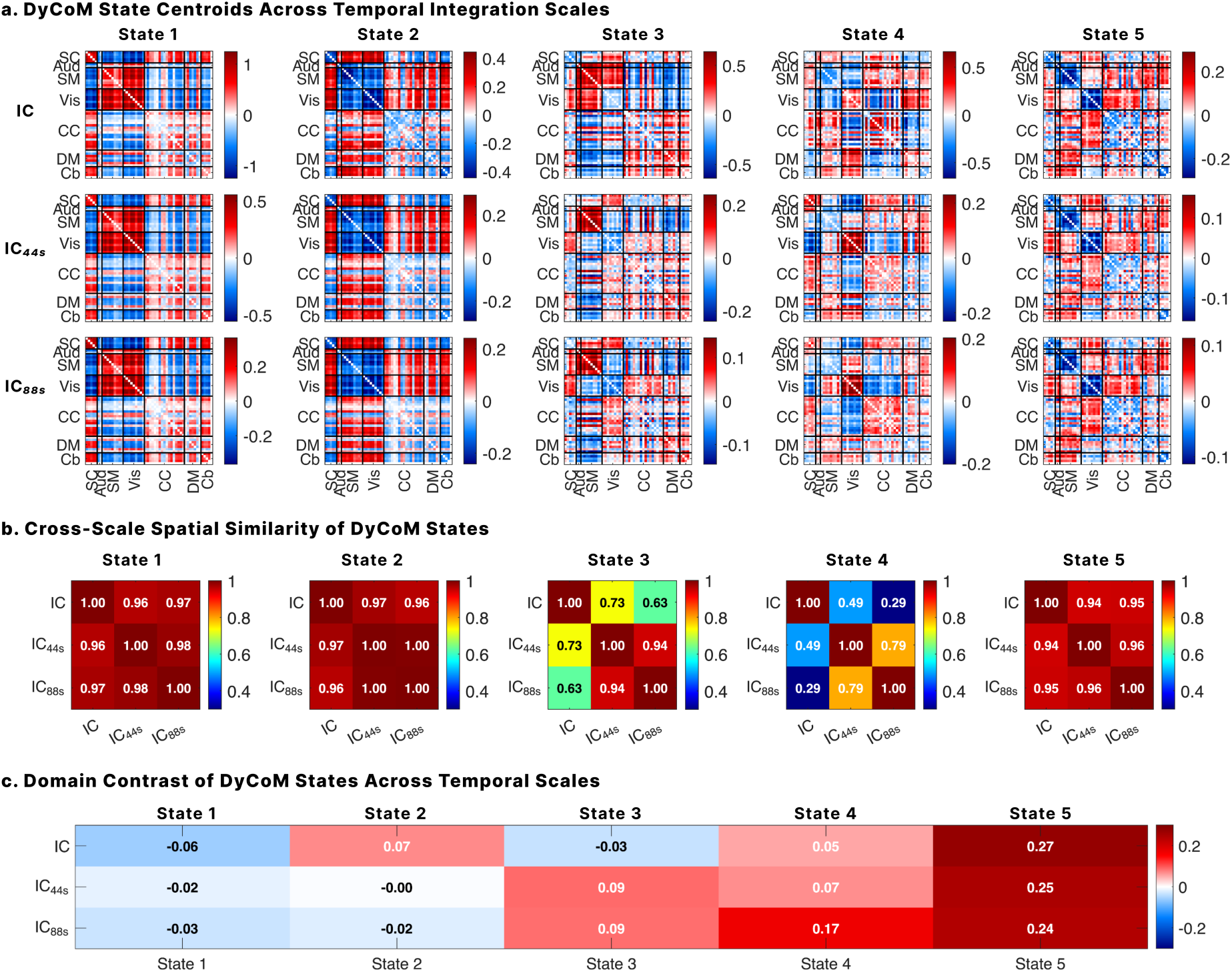
Instantaneous and windowed DyCoM formulations reveal scale-dependent state structure. **a.** State centroids estimated using instantaneous correlation (IC), 44 s integration (IC_44s_), and 88 s integration (IC_88s_), with all operators held constant except the temporal kernel. States 1, 2, and 5 exhibit strong visual similarity across integration scales, whereas states 3 and 4 show clear scale-dependent reconfiguration. **b.** Cross-scale spatial similarity between state centroids, quantified using Pearson correlation. States 1, 2, and 5 demonstrate high similarity across scales, while states 3 and 4 display reduced similarity, particularly between instantaneous and longer integration conditions. **c.** Domain segregation contrast across temporal scales. Temporal integration increases within-domain dominance for states 3 and 4 relative to instantaneous interaction, whereas states 1 and 2 remain near zero across scales. These results illustrate that DyCoM accommodates both instantaneous and windowed formulations within a unified operator structure while revealing that certain states are intrinsically scale

## 4. DISCUSSION

### 4.1. DyCoM as a unifying operator framework for time-varying connectivity

This work introduces dynamic co-modulation as a unified operator framework for dynamic connectivity analysis. Rather than focusing on a specific estimator or data modality, DyCoM defines interaction through a sequence of modular operations: representation, instantaneous interaction, temporal integration, and normalization. This decomposition exposes the shared computational structure underlying widely used correlation-based dynamic connectivity methods, including instantaneous correlation, sliding window correlation, and adaptive formulations, and clarifies how differences between methods arise from specific operator choices rather than from fundamentally distinct analytical principles. By separating estimator logic from empirical interpretation, DyCoM promotes coherence across findings, improves interpretability, and supports principled methodological development in studies of evolving system interactions. Crucially, this operator-level separation enables distinction between reproducibility driven by true system dynamics and apparent agreement arising from shared estimator structure.

Crucially, DyCoM is grounded in classical signal processing theory. The instantaneous interaction layer corresponds to bilinear energy mappings, which have long been studied in adaptive and time-frequency signal analysis. The representation and normalization stages mirror standard operations used to stabilize nonstationary signals under noise and drift. By framing dynamic connectivity as a form of signal co-modulation, DyCoM connects contemporary neuroimaging estimators to well-established principles from communication theory and time-frequency analysis^23^. This grounding provides a principled basis for interpreting how representation, integration, and normalization choices shape dynamic connectivity estimates, and it enables new estimators to be constructed systematically rather than introduced as ad hoc methodological variants. Although motivated by neuroimaging, the DyCoM operators correspond to generic signal processing constructs that apply to any multivariate time series in which interactions are inferred from co-modulated activity.

### 4.2. Robustness to nuisance as a function of operator composition

The simulation results illustrate how robustness to nuisance components emerges from the composition of operators within the DyCoM framework rather than from any single estimator in isolation. Under clean conditions, all parameterizations accurately recovered the prescribed connectivity trajectory, confirming that the estimators are unbiased in the absence of confounds. As nuisance components were introduced, differences across methods appeared in a structured manner that directly reflected which DyCoM operators were active.

When nuisance effects were weak and shared across regions, parameterizations relying solely on instantaneous interaction were more susceptible to contamination, whereas incorporating temporal integration and interaction-level normalization reduced sensitivity to shared low frequency variance. As nuisance strength increased, particularly under conditions of strong or asymmetric contamination, methods lacking adaptive representation showed marked degradation, while parameterizations including local representation remained comparatively stable. In this regime, normalization at the interaction level further constrained variability without altering the underlying state structure.

Viewed through the DyCoM framework, these patterns clarify why different estimators perform similarly under ideal conditions yet diverge systematically as nuisance severity increases. Robustness is not an intrinsic property of a specific method, but instead reflects how representation, interaction, integration, and normalization operators jointly shape sensitivity to shared and unshared confounds. This operator-level perspective replaces heuristic distinctions between methods with a mechanistic understanding of how robustness arises.

### 4.3. Complementary dynamic signatures across DyCoM parameterizations

Application of DyCoM to resting-state fMRI revealed that different parameterizations emphasize distinct and complementary aspects of schizophrenia-related dysconnectivity. These differences are not merely computational but reflect known neurobiological signatures of the disorder. For example, the dominant state identified by IC was characterized by a fragmented profile with positive weights in higher-order visual regions (e.g., cuneus and lingual gyrus) and negative weights in primary sensory and parietal areas. This pattern aligns with prior studies reporting increased engagement of visual and sensorimotor networks in schizophrenia, particularly in dynamic states linked to perceptual overload or hyper-responsivity^47,48^. IC’s sensitivity to these components suggests that minimally normalized bilinear interactions may be well suited to detect sensory-driven disruptions.

SWPC, which adds moment-based normalization to the IC framework, captured a smoother posterior–anterior gradient, with strong contributions from visual and temporal cortices and negative weights in frontal and parietal regions. This spatial pattern mirrors previous findings of imbalances between fronto-temporal and sensorimotor networks in schizophrenia ^48,49^. Although SWPC exhibited lower overall diagnosis sensitivity across metrics, its smooth network gradients and longer temporal scale suggest it captures slower, more stable connectivity shifts relevant to broader macroscale reorganization.

Both aIC and saIC produced highly similar dominant states, characterized by strong positive loadings in visual, parietal, and sensorimotor regions and negative loadings in frontal and subcortical areas. This network configuration overlaps with disruptions in the default mode and executive control systems that have been implicated in schizophrenia symptoms and cognitive deficits^14,50,51^. The adaptive representation in aIC and the additional moment-based normalization in saIC may enhance sensitivity to these broad patterns by suppressing shared variance and improving signal-to-noise at both the representation and interaction levels. Notably, these adaptive methods yielded consistent dominant states across all three dynamic metrics, suggesting greater stability in state definition across analytic resolutions.

In terms of statistical sensitivity, IC showed the strongest group differences for mean dwell time and transition matrix statistics, suggesting it is especially sensitive to state stability and switching behavior. By contrast, aIC and saIC demonstrated the strongest effects for fractional occupancy (FR), indicating a heightened sensitivity to state prevalence and global co-fluctuation structure. These distinctions further reflect the computational design of each method: while IC responds directly to raw co-modulation, aIC and saIC emphasize localized fluctuations that persist across time, capturing different temporal properties of brain state organization.

Associations with clinical symptoms further reinforce these distinctions. IC and SWPC metrics were most strongly associated with negative symptom severity, consistent with the idea that traditional correlation-based approaches preferentially detect hypoactive or rigid network patterns^52,53^. In contrast, aIC and saIC exhibited more balanced associations with both symptom domains. While IC and SWPC exhibited minimal correspondence with antipsychotic dose (CPZ), aIC and saIC showed stronger CPZ associations. This enhanced sensitivity to CPZ may arise from the adaptive representation layer, which performs local z-scoring, thereby suppressing slow amplitude fluctuations (i.e., gain). This process reduces amplitude dominance and preserves timing structure, aligning with prior findings that medication effects are more strongly associated with timing-based features than with absolute co-activation magnitude^42,43,54^. The result is a co-modulation profile that leans toward the fluctuation-timing regime, potentially enhancing sensitivity to treatment-related dynamics. In this view, the CPZ coupling of the adaptive estimators is not merely a medication confound but a potential window into treatment-related reorganization of network dynamics. These medication-related associations should nevertheless be interpreted with caution, as illness chronicity and treatment duration were not available in the present dataset and could not be included in the covariate structure.

These distinctions help explain why prior studies have implicated divergent networks, from visual hyper-responsivity to executive control deficits, depending on analytic choices. At the same time, they clarify why multiple estimators can yield overlapping behavioral and clinical association profiles, including similar strengths of association with cognitive measures, without implying methodological equivalence or biological redundancy. DyCoM reveals both divergence and apparent agreement as consequences of specific operator configurations, showing that parameterizations represent orthogonal projections of the brain’s dynamic organization. By clarifying how estimator design governs sensitivity to neural and clinical features, DyCoM provides a principled foundation for interpreting dynamic connectivity findings and supports complementary, interpretable use of multiple parameterizations in clinical neuroscience.

The simulations also delineate when each family is preferable. Non-adaptive estimators are the more accurate choice when the coupled neural fluctuations are slow and structured confounds are controlled or spectrally distant (Scenario 5); adaptive estimators are preferable when a spectral margin separates confound from coupling (Scenarios 2 to 4 and S7), and no estimator succeeds when that margin vanishes (Scenario 6). Because unstructured noise degrades all parameterizations equally (Scenario S9), the method differences observed in the fMRI analyses reflect structured properties of the data interacting with operator choices rather than differential noise sensitivity.

### 4.4. Temporal integration as an explicit modeling dimension

An important implication of the operator formulation is that temporal integration is not a required feature of dynamic connectivity estimation, but a configurable component within a broader computational structure. Instantaneous and temporally integrated connectivity are simply distinct instantiations of the same operator sequence. By varying only the temporal integration operator, we observed that some states were stable across scales, whereas others exhibited pronounced scale dependence. Temporal integration enhanced within-domain dominance for specific states that appeared weakly organized in the instantaneous formulation. This pattern indicates that sustained, domain-coherent interactions become more prominent with temporal accumulation, whereas instantaneous formulations emphasize rapid, distributed co-fluctuations across brain networks. Rather than assuming that windowing defines dynamic connectivity, DyCoM makes temporal integration an explicit modeling decision, allowing investigators to choose the temporal regime appropriate for their scientific question while preserving a unified mathematical framework.

### 4.5. Scope, limitations, and extensions

The present study focuses on amplitude-based co-modulation, corresponding to correlation-based dynamic connectivity, due to its widespread use and methodological prominence in fMRI research. While the DyCoM framework naturally extends to phase-and frequency-based co-modulation via alternative representation and interaction operators, such instantiations were not empirically evaluated here. Similarly, DyCoM is not intended to subsume causal or directed connectivity models, which pursue distinct inferential aims that fall outside the scope of this work. More generally, DyCoM unifies the family of estimators that quantify co-modulation from observed signals; the present study instantiates and validates amplitude-based parameterizations, though the operator structure extends in principle to phase- and frequency-based co-modulation. Approaches such as switching hidden Markov models, change-point detection, or dimensionality-reduction summaries operate at a different level of the analysis pipeline: they summarize already-estimated connectivity features and can be applied downstream of any DyCoM parameterization, complementing rather than competing with the framework. Likewise, directed and generative models, including Granger causality, transfer entropy, dynamic causal modelling, and state-space formulations, characterize the processes generating the data rather than the observed covariance structure, and remain outside the present scope.

Nevertheless, the operator structure of DyCoM provides a foundation for future extensions. These may include phase synchrony and cross-frequency coupling estimators, alternative nonlinear or kernelized energy mappings, or online adaptations for real-time applications. Importantly, our findings show that even within the widely used correlation-based regime, substantial methodological diversity can be traced to a small set of interpretable operator choices, enabling compact expression, systematic extension, and principled evaluation.

## 5. Author contribution

**Sir-Lord Wiafe:** Conceptualization, Formal analysis, Methodology, Visualization, Writing –original draft. **Najme Soleimani:** Formal analysis, Writing – review & editing. **Armin Iraji:** Writing – review & editing. **Tülay Adali:** Writing – review & editing. **Vince Calhoun:** Conceptualization, Funding acquisition, Validation, Methodology, Resources, Supervision, Writing –original draft, Writing – review & editing.

## 8. DATA & CODE AVAILABILITY STATEMENT

The data was not collected by us and was provided in a deidentified manner. The IRB will not allow sharing of data or individual derivatives, as a data reuse agreement was not signed by the subjects during the original acquisition. The DyCoM framework is fully specified mathematically in the Methods and Supplementary Material, sufficient for independent implementation. The MATLAB implementation is available from the corresponding author upon reasonable request; a public release is planned following completion of a pending intellectual property filing.

