## supplementary material for "Dynamic Co-Modulation (DyCoM): A Unified Operator Framework for Dynamic Connectivity in Neuroimaging"

### *Supplementary Section S1. Relation between the DyCoM Bilinear Kernel and the Cross Wigner–Ville Distribution*

#### S1.1 Objective

This section demonstrates that the instantaneous bilinear energy kernel used in the DyCoM framework,

$$\Phi_{ij}(t;\mathcal{P}) = \tilde{x}_i(t) \cdot \tilde{x}_j^*(t) \tag{S1}$$

is mathematically equivalent to the zero-lag component of the classical cross Wigner–Ville distribution (XWVD) between signals $\tilde{x}_i(t)$ and $\tilde{x}_j(t)$.

For real-valued signals, conjugation reduces to the identity, yielding $\Phi_{ij} = \tilde{x}_i(t) \cdot \tilde{x}_j(t)$, as used in the main text.

#### S1.2 Definition of the cross Wigner–Ville distribution

For two analytic (possibly complex) signals $\tilde{x}_i(t)$ and $\tilde{x}_j(t)$ the cross Wigner–Ville distribution is the Fourier transform of their lagged product:

$$W_{ij}(t,f) = \int_{-\infty}^{\infty} \tilde{x}_i\left(t+\frac{\tau}{2}\right) \cdot \tilde{x}_j^*\left(t-\frac{\tau}{2}\right) e^{-j2\pi f\tau} d\tau \tag{S2}$$

#### S1.3 Integrating the XWVD over all frequencies

Integrating $W_{ij}(t,f)$ over frequency yields:

$$\int_{-\infty}^{\infty} W_{ij}(t,f)df = \int_{-\infty}^{\infty}\left[\int_{-\infty}^{\infty} \tilde{x}_i\left(t+\frac{\tau}{2}\right) \cdot \tilde{x}_j^*\left(t-\frac{\tau}{2}\right) e^{-j2\pi f\tau} d\tau\right] df$$

$$= \int_{-\infty}^{\infty} \tilde{x}_i\left(t+\frac{\tau}{2}\right) \cdot \tilde{x}_j^*\left(t-\frac{\tau}{2}\right)\left[\int_{-\infty}^{\infty} e^{-j2\pi f\tau}\, df\right] d\tau$$

Exchanging the order of integration and applying the Fourier identity (S3)

$$\int_{-\infty}^{\infty} e^{-j2\pi f\tau}\, df = \delta(\tau),$$

gives

$$\int_{-\infty}^{\infty} W_{ij}(t,f)df = \int_{-\infty}^{\infty} \tilde{x}_i\left(t+\frac{\tau}{2}\right) \cdot \tilde{x}_j^*\left(t-\frac{\tau}{2}\right) \delta(\tau) d\tau$$

At $\tau = 0$ (zero-lag), the delta collapses the $\tau$-integration:

$$\int_{-\infty}^{\infty} W_{ij}(t,f)df = \tilde{x}_i(t)\tilde{x}_j^*(t) = \Phi_{ij}(t) \quad \text{(S4)}$$

#### S1.4 Interpretation

The instantaneous DyCoM kernel, therefore, corresponds to the zero-lag projection of the full bilinear time–frequency representation:

- It captures the instantaneous cross-energy shared between two signals at time $t$.
- Integrating $W_{ij}(t,f)$ over frequency summarizes spectral information but preserves local covariance structure.
- In DyCoM, subsequent timescale integration via the kernel, $h(\tau)$, generalizes this instantaneous bilinear energy into adaptive, multi-timescale co-modulation estimates.

Thus, the instantaneous term

$$\Phi_{ij}(t) = \tilde{x}_i(t)\tilde{x}_j^*(t)$$

used in DyCoM is not heuristic but the theoretically exact zero-lag limit of the cross Wigner–Ville distribution, linking dynamic co-modulation to the broader family of **bilinear** time–frequency energy operators.

### *Supplementary Section S2. saIC approaches SWPC as a limiting case*

Let $h_T(\tau)$ be a normalized temporal kernel (sliding window) with $\sum_\tau h_T(\tau) = 1$.

**SWPC.** Define the windowed moments with a window length $T$

$$\mu_i^{(T)}(t) = \sum_\tau h_T(\tau)\, x_i(t-\tau), \qquad (\sigma_i^{(T)}(t))^2 = \sum_\tau h_T(\tau)(x_i(t-\tau) - \mu_i^{(T)}(t))^2,$$

and similarly, for $j$. The sliding-window Pearson correlation is

$$\mathrm{SWPC}_{ij}(t) = \frac{\sum_\tau h_T(\tau)\left(x_i(t-\tau) - \mu_i^{(T)}(t)\right)\left(x_j(t-\tau) - \mu_j^{(T)}(t)\right)}{\sigma_i^{(T)}(t)\,\sigma_j^{(T)}(t)}. \quad \text{(S5)}$$

**saIC (DyCoM form).** Let $h_R(\tau)$ be a representational kernel with window length $R$ and define

$$\mu_i^{(R)}(t) = \sum_\tau h_R(\tau)\, x_i(t-\tau), \qquad (\sigma_i^{(R)}(t))^2 = \sum_\tau h_R(\tau)(x_i(t-\tau) - \mu_i^{(R)}(t))^2,$$

with analogous definitions for $j$. Define the standardized representations

$$\tilde{x}_i(t) = \frac{x_i(t) - \mu_i^{(R)}(t)}{\sigma_i^{(R)}(t)}, \qquad \tilde{x}_j(t) = \frac{x_j(t) - \mu_j^{(R)}(t)}{\sigma_j^{(R)}(t)}. \qquad \text{(S6)}$$

The instantaneous interaction is

$$\Phi_{ij}(t) = \tilde{x}_i(t)\tilde{x}_j(t), \qquad \text{(S7)}$$

and the temporally integrated interaction is

$$S_{ij}(t) = \sum_{\tau} h_T(\tau)\, \Phi_{ij}(t-\tau) = \sum_{\tau} h_T(\tau)\, \tilde{x}_i(t-\tau)\tilde{x}_j(t-\tau). \qquad \text{(S8)}$$

**Expansion.** Substitute (S6) into (S7):

$$saIC_{ij}(t) = \sum_{\tau} h_T(\tau)\, \frac{x_i(t-\tau) - \mu_i^{(R)}(t-\tau)}{\sigma_i^{(R)}(t-\tau)} \cdot \frac{x_j(t-\tau) - \mu_j^{(R)}(t-\tau)}{\sigma_j^{(R)}(t-\tau)}. \qquad \text{(S9)}$$

**Limiting regime $R \gg T$.** When $R$ is large relative to $T$, the representational moments are approximately constant over the temporal window, and they serve the same role as the windowed local moments used by SWPC. Therefore over the support of $h_T$,

$$\mu_i^{(R)}(t-\tau) \approx \mu_i^{(R)}(t), \qquad \sigma_i^{(R)}(t-\tau) \approx \sigma_i^{(R)}(t), \qquad \text{(S10)}$$

and similarly, for $j$. Applying (S10) in (S9) yields

$$S_{ij}(t) \approx \sum_{\tau} h_T(\tau)\, \frac{x_i(t-\tau) - \mu_i^{(R)}(t)}{\sigma_i^{(R)}(t)} \cdot \frac{x_j(t-\tau) - \mu_j^{(R)}(t)}{\sigma_j^{(R)}(t)}. \qquad \text{(S11)}$$

Factor out the time-$t$ denominators:

$$S_{ij}(t) \approx \frac{1}{\sigma_i^{(R)}(t)\sigma_j^{(R)}(t)} \sum_{\tau} h_T(\tau)\, \left(x_i(t-\tau) - \mu_i^{(R)}(t)\right)\left(x_j(t-\tau) - \mu_j^{(R)}(t)\right). \qquad \text{(S12)}$$

**Identification with SWPC.** Under the same regime, the representational moments are effectively constant within the temporal window and serve as the window-local centering and scaling terms. Thus, identifying $\mu_i^{(R)}(t) \approx \mu_i^{(T)}(t)$ and $\sigma_i^{(R)}(t) \approx \sigma_i^{(T)}(t)$ gives

$$S_{ij}(t) \approx \mathrm{SWPC}_{ij}(t). \qquad \text{(S13)}$$

Therefore, when the representational window varies slowly relative to the temporal integration window ($R \gg T$), the saIC temporally integrated interaction reduces to the classical sliding window Pearson correlation as a limiting case.

Note that the derivation above concerns the temporally integrated interaction, $S_{ij}(t)$, while the full saIC additionally applies moment-based normalization computed over the same temporal kernel. In the R ≫ T regime, the standardized representations differ from the raw signals only by centering and scaling terms that are effectively constant within the temporal window, and because the Pearson correlation is invariant to such locally affine transformations, applying the final normalization leaves the identification with SWPC unchanged.

### Supplementary Section S3. Additional simulation scenarios (S7–S10)

To further evaluate robustness beyond the scenarios reported in the main text, four additional simulations were constructed using the same generative model described in Section 2.4 of the main manuscript. Unless stated otherwise, two regional time courses were simulated at fs = 1 Hz for T = 2000 s, with the ground-truth connectivity trajectory $\rho(t)$ generated by band-limiting Gaussian white noise to 0.002–0.004 Hz and scaling its amplitude to a maximum of 0.9; the neural sources were filtered to the canonical BOLD band (0.01–0.15 Hz) and mixed through a time-dependent covariance matrix so that their instantaneous correlation matched $\rho(t)$. All four estimators shared the same Hamming FIR temporal integration (88 s), with the representational window for aIC and saIC set to 33 s. Each scenario comprised 1000 independent trials with a fixed ground-truth trajectory; all time series were z-scored before analysis, and performance was evaluated over t = 300–1700 s to exclude filter and window edge effects.

*(i) Multiple simultaneous nuisance sources (S7).* Whereas Scenarios S2–S4 examined one confound class at a time, real resting-state data contain multiple temporally overlapping confounds with distinct spectral and spatial profiles; this scenario tests whether the robustness observed under isolated contamination generalizes to compound contamination. Four confounds were added concurrently: a shared scanner-drift component band-limited to 0.001–0.005 Hz (weight 0.6 on both nodes); a shared respiratory/$CO_2$-like component band-limited to 0.02–0.04 Hz with region-specific susceptibility (weights 0.8 and 1.0); region-specific vascular components band-limited to 0.05–0.08 Hz, generated independently per node (weight 0.5); and motion-like transient bursts, modeled as sparse impulses at approximately 2% of time points with amplitudes drawn from $3 \cdot \mathcal{N}(0,1)$, smoothed with a 9 s Hann kernel and generated independently per node. All nuisance components were standardized to unit variance before weighting.

*(ii) Abrupt connectivity transition (S8).* Because the main simulations prescribe smooth, slowly varying coupling, they cannot reveal whether any parameterization smears or destabilizes around discontinuous state changes associated with task onsets and spontaneous state switching; this scenario directly tests transition tracking. The ground-truth trajectory was replaced by a step function, $\rho(t) = 0.15$ for $t < 1000$ s and $\rho(t) = 0.75$ thereafter, with no nuisance contamination. The temporal kernel was set to 44 s, matching

the window length used in the fMRI analyses, to assess transition tracking at the empirical timescale.

*(iii) Low signal-to-noise ratio (S9).* This scenario serves as a control that separates sensitivity to unstructured noise from sensitivity to structured nuisance: if the parameterizations degraded unequally under white noise alone, the method differences observed in Scenarios S2–S4 and S7 could reflect robustness to generic noise rather than the interaction between structured confounds and specific operator choices. Independent white Gaussian measurement noise of variance equal to the signal variance (SNR = 1) was added to each node, with no structured confound.

*(iv) Fast time-varying connectivity with matched integration (S10).* Because the main simulations examine coupling in the 0.002–0.004 Hz range where windowed estimators are expected to perform well, this scenario tests whether faster coupling dynamics remain recoverable, and whether the accessible timescale is set by the temporal integration operator rather than by estimator identity (Section 4.4). The ground-truth trajectory was band-limited to 0.01–0.02 Hz (periods of 50–100 s), and the temporal kernel was shortened to 22 s so that the integration timescale matched the coupling dynamics. No nuisance was added.

As in the main simulations, estimated trajectories were compared to the ground truth using Pearson correlation and root-mean-square error, with distributions summarized across the 1000 trials (Supplementary Figure S1).

### Figure S1: Robustness of DyCoM parameterizations across additional simulation scenarios (S7–S10)

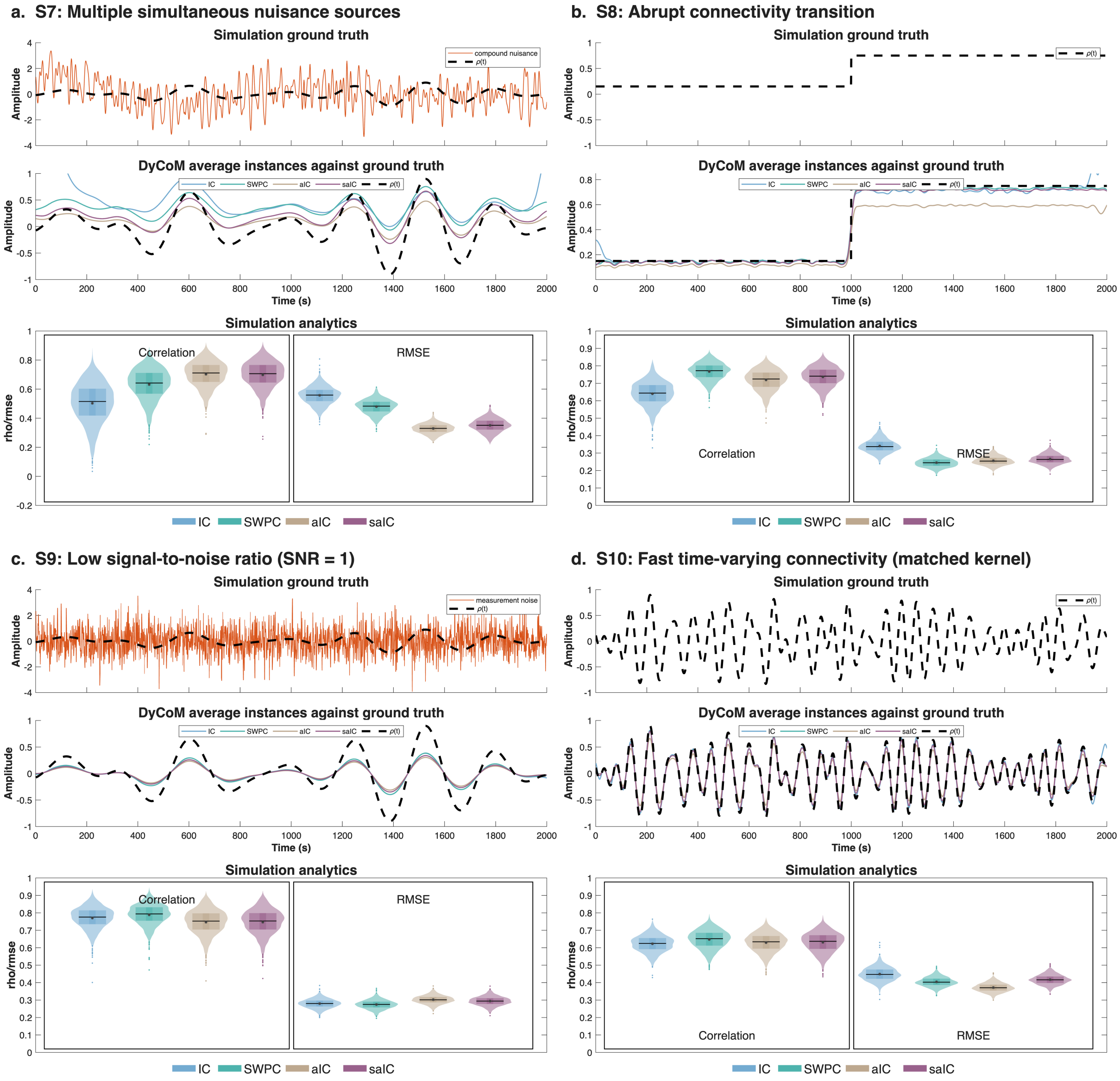


*Simulation scenarios evaluate the four DyCoM parameterizations, IC, SWPC, aIC, and saIC, under compound, non-stationary, noisy, and fast-coupling regimes. In each panel, the top row shows the ground truth connectivity trajectory (ρ(t)) with or without simulated noise. The middle row plots average estimated trajectories across 1000 trials, overlaid with the ground truth. The bottom row presents violin plots of estimator performance, quantified by Pearson correlation and root-mean-square error (RMSE) relative to ρ(t).* ***a.*** *Scenario S7 combines multiple simultaneous nuisance sources, where aIC and saIC remain the most accurate.* ***b.*** *In Scenario S8, an abrupt connectivity transition is tracked by all estimators without instability.* ***c.*** *Scenario S9 adds strong unstructured measurement noise, which degrades all estimators uniformly.* ***d.*** *Scenario S10 prescribes faster coupling dynamics with a matched temporal kernel, which all estimators recover comparably. These results indicate that performance differences among estimators arise from structured nuisance interacting with specific operator choices, rather than from general noise sensitivity or timescale limitations.*

**Figure S2: Sensitivity of adaptive DyCoM parameterizations to window-length choices**

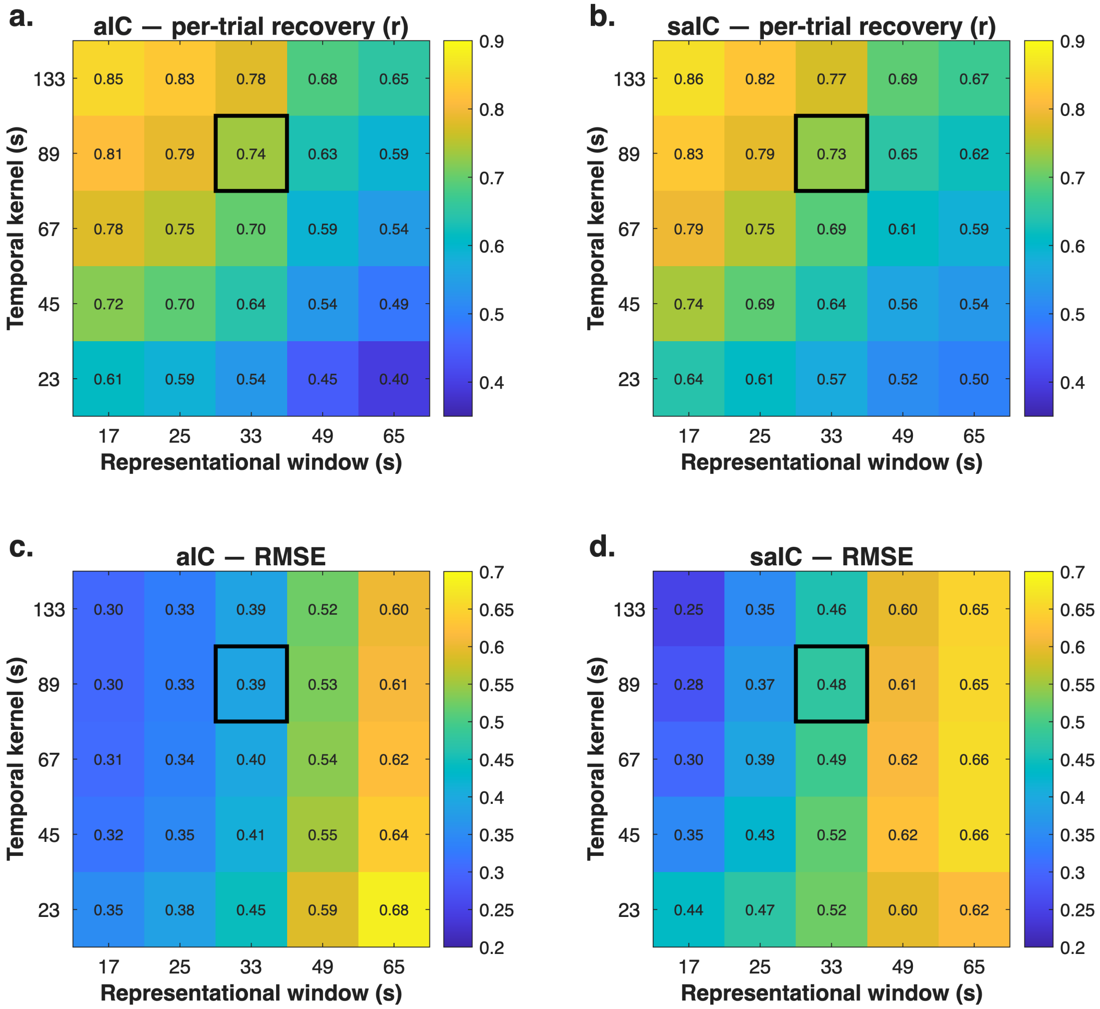


*Systematic parameter sweep over the temporal integration kernel (22–132 s) and the representational window (17–65 s) for aIC and saIC under the strong shared nuisance scenario (Scenario 3), with values averaged across 100 trials. The top row shows per-trial recovery (Pearson correlation with ρ(t)), and the bottom row shows RMSE; the black outline marks the settings used in the main simulations (88 s integration, 33 s representation).* ***a, b.*** *Recovery varies smoothly across the grid, improving with longer integration kernels and shorter representational windows.* ***c, d.*** *RMSE shows the complementary gradient, increasing as the representational window lengthens and its cutoff falls below the nuisance band. Performance changes gradually across all settings, with no abrupt sensitivity to either parameter, and the gradients follow the expected operator behavior: the temporal kernel governs averaging of the slowly varying coupling, while the representational window governs nuisance suppression. For reference, SWPC evaluated at the same integration kernels yields r = 0.50–0.65 and RMSE = 0.63–0.69, which the adaptive parameterizations exceed across the recommended range and approach only when the representational window is deliberately mis-set.*

**Table S 1 | Numerical equivalence between DyCoM parameterizations and standard implementations**

| DyCoM Parameterization | Independent standard implementation | Max \|difference\| |
|---|---|---|
| SWPC (rectangular window) | Per-window Pearson correlation | $2.1 \times 10^{-8}$ |
| SWPC (Hamming window) | Weighted Pearson correlation | $1.6 \times 10^{-8}$ |
| IC | Edge time series $z_i(t) \cdot z_j(t)$ | 0 (exact) |
| Exponential (EWMA) kernel | Recursion $y_t = \alpha y_{t-1} + (1-\alpha) x_t$ | 0 (exact) |
| FBC (aIC + filter bank) | Local z (movmean/movstd) + Butterworth filter | $1.4 \times 10^{-14}$ |

Each DyCoM parameterization was applied to the same simulated time series (T = 1200 s) as an independently implemented standard method, and the maximum absolute difference between the two outputs was computed across all time points. Differences at the $10^{-8}$ level reflect the $\varepsilon = 10^{-8}$ variance regularization used in the DyCoM implementation; all remaining differences are at floating-point precision. These results complement the analytical correspondences in Supplementary Sections S1–S2 by confirming that the DyCoM implementations reproduce widely used estimators exactly, so that results obtained within the framework are directly comparable to prior literature.

### Figure S3: Cluster-number selection across DyCoM parameterizations

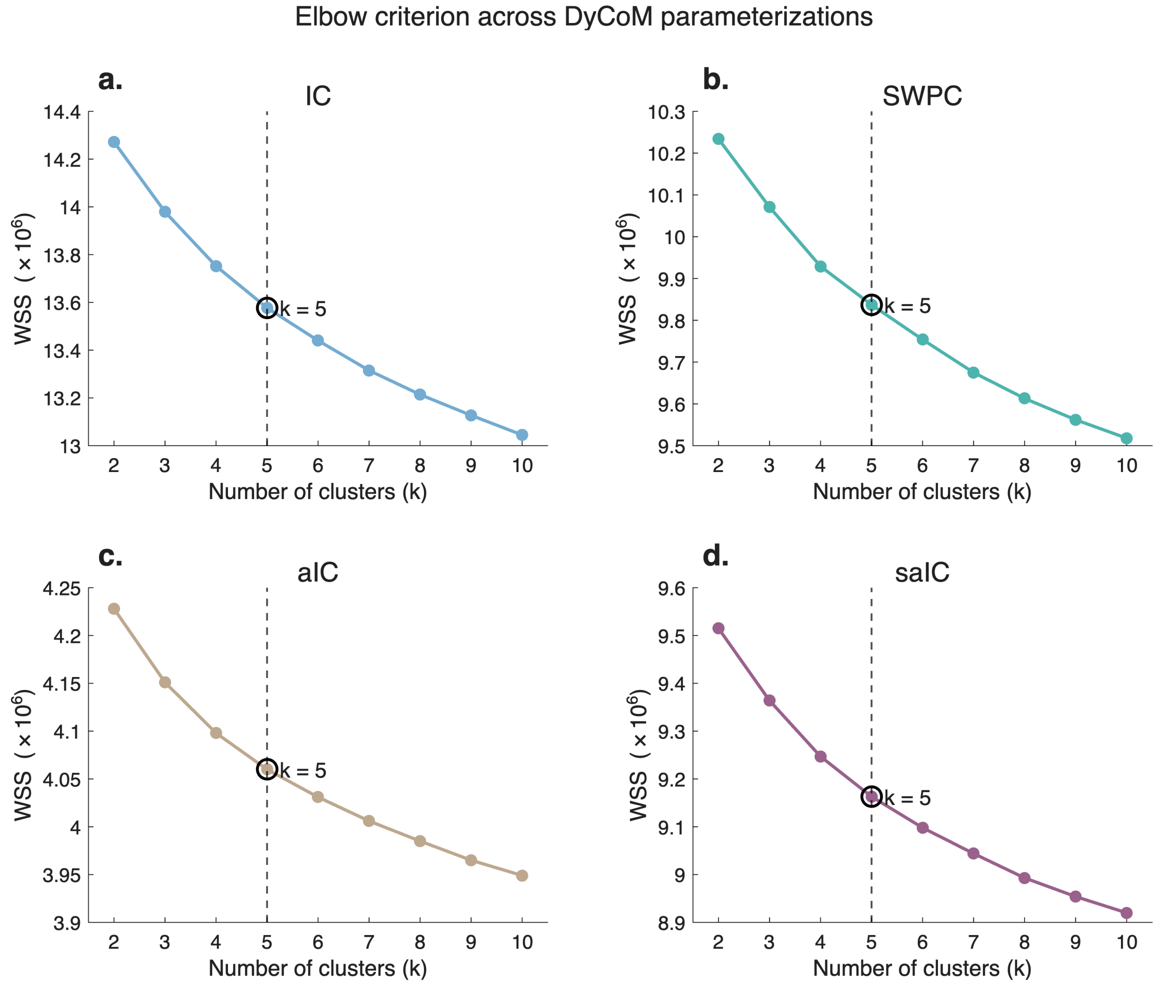


*Elbow curves of the within-cluster sum of squares (WSS) from k-means clustering, computed separately for each DyCoM parameterization over k = 2 to 10 (a single-cluster solution does not constitute a meaningful state decomposition). The optimal number of clusters, identified with the kneedle criterion as the point of maximum deviation from the chord connecting the curve endpoints, is k = 5 for all four parameterizations (dashed line, circled).* ***a.*** *IC.* ***b.*** *SWPC.* ***c.*** *aIC.* ***d.*** *saIC. The agreement across parameterizations indicates that the five-state decomposition used in the main analyses reflects a shared optimum rather than an imposed common solution.*
